# An Ionic Sensor acts in Parallel to dSarm to Promote Neurodegeneration

**DOI:** 10.1101/2024.10.29.620922

**Authors:** Adel Avetisyan, Romina Barria, Amy Sheehan, Marc R. Freeman

## Abstract

How neurons to sense when they are terminally dysfunctional and activate neurodegeneration remains poorly defined. The pro-degenerative NAD^+^ hydrolase dSarm/SARM1 can act as a metabolic sensor by detecting pathological changes in NAD^+^/NMN and subsequently induce catastrophic axon degeneration. Here we show *Drosophila* with-no-lysine kinase (dWnk), which can directly sense Cl^-^, K^+^ and osmotic pressure, is required for neurodegeneration induced by depletion of the NAD^+^ biosynthetic enzyme dNmnat. dWnk functions in parallel to dSarm and acts through the downstream kinase Frayed to promote axon degeneration and neuronal cell death. dWnk and dSarm ultimately converge on the BTB-Back domain molecule Axundead (Axed) to execute neurodegeneration. Our work argues that neurons use direct sensors of both metabolism (dSarm/SARM1) and ionic/osmotic status (dWnk) to evaluate cellular health and, when dysfunctional, promote neurodegeneration though a common axon death signaling molecule, Axundead.

## Introduction

Neurons are highly metabolically active and morphologically complex cells, and as such they are particularly vulnerable to a variety of insults, including metabolic disorders, mitochondrial dysfunction, toxin exposure, and physical injury. These vulnerabilities can lead to neurodegeneration and the loss of neuronal connectivity, hallmark features of neurological diseases such as Alzheimer’s and Parkinson’s^1–4^, but how neurodegeneration is triggered in any context remains poorly understood^4^. Studies beginning with the *Wallerian degeneration slow* (*Wld^S^*) molecule^5^ revealed that axonal survival critically depends on the delivery of the NAD^+^ biosynthetic enzyme NMNAT2 from the cell body to the axon^6,7^. After axon injury, delivery of NMNAT2 is blocked, the labile pool of axonal NMNAT2 molecule is degraded^8^, NAD^+^ and ATP are depleted, and the NMNAT2 substrate NMN accumulates. This in turn binds and activates the pro-degenerative metabolic sensor dSarm/SARM1 (*sterile alpha/Armadillo/Toll-Interleukin receptor (TIR) homology domain protein*) by binding to its ARM domain^9–13^ and disinhibiting its intrinsic NAD^+^ hydrolase activity, thereby driving further catastrophic depletion of NAD^+^ and neurodegeneration^14,15^. Remarkably, while *NMNAT2* null mouse mutants die perinatally with short axons, *NMNAT2, SARM1* double mutants survive and appear phenotypically normal, arguing that a significant part of the pro-degenerative signaling that occurs after NMNAT2 loss flows through SARM1^16^.

In *Drosophila*, dSarm activation promotes neurodegeneration through a BTB and C-terminal Kelch (BACK) domain protein Axundead (Axed). Loss of Axed is sufficient to completely block the degenerative effects of dSarm activation^17^, and *dSarm* and *axed* mutants are also capable of suppressing neurodegeneration induced by partial depletion of *Drosophila* Nmnat (dNmnat) by RNAi^17^, indicating that depletion of NMNAT molecules is a conserved mechanism for SARM1/dSarm activation. Surprisingly, while *dSarm* mutants can suppress neurodegeneration induced by partial depletion of dNmnat, *dSarm* null mutants cannot rescue *dNmnat* null mutant clones, although *axed* mutants can^17^. This indicates that *axed* mutants are even more neuroprotective than *dSarm* mutants and argues that additional signaling mechanisms parallel to dSarm must exist to drive neurodegeneration^17^. Here we identify the ionic and osmotic sensor *Drosophila* with-no-lysine kinase (dWnk) and its downstream target kinase Frayed as pro-degenerative signaling molecules. After dNmnat depletion, dWnk and Frayed act in parallel to dSarm, ultimately converging on Axed, to promote neurodegeneration. We propose these parallel sensors for ionic/osmotic (dWnk) and metabolic status (dSarm/SARM1) continuously sense intrinsic neuronal health and that can trigger neurodegeneration in pathologically compromised neurons.

## Results

### *Drosophila* With-no-lysine kinase (dWnk) promotes neurodegeneration induced by dNmnat depletion

To identify new molecules that drive neurodegeneration after dNmnat depletion, we performed an unbiased EMS-based forward genetic screen for mutations that cell-autonomously suppressed neurodegeneration in the presence of dNmnat^RNAi^. We labeled glutamatergic sensory neurons in the L1 nerve of the *Drosophila* anterior wing vein, which extend their axons medially into the thoracic nervous system^18,19^. Mosaic Analysis with a Repressible Cell Marker (MARCM) was used to simultaneously label single-cell clones with membrane-tethered GFP and induce neurodegeneration by depleting dNmnat using a *UAS-RNAi* construct (Fig. S1A)^18,20^. In control MARCM clones (no RNAi), the majority of glutamatergic sensory neuron cell bodies, dendrites, and axons remained intact for the lifespan of the fly. However, when we drove a *UAS-dNmnat^RNAi^* construct in MARCM clones, we found that on average half of the labeled axons showed dying back degeneration already at 1 day post eclosion (dpe) while the majority of the cell bodies and dendrites remained morphologically intact (Figure S1A, D, E). By 10 dpe, >85% axons and dendrites were cleared along with most neuronal cell corpses (Fig 1B-D and S1D, E).

**Figure 1.**
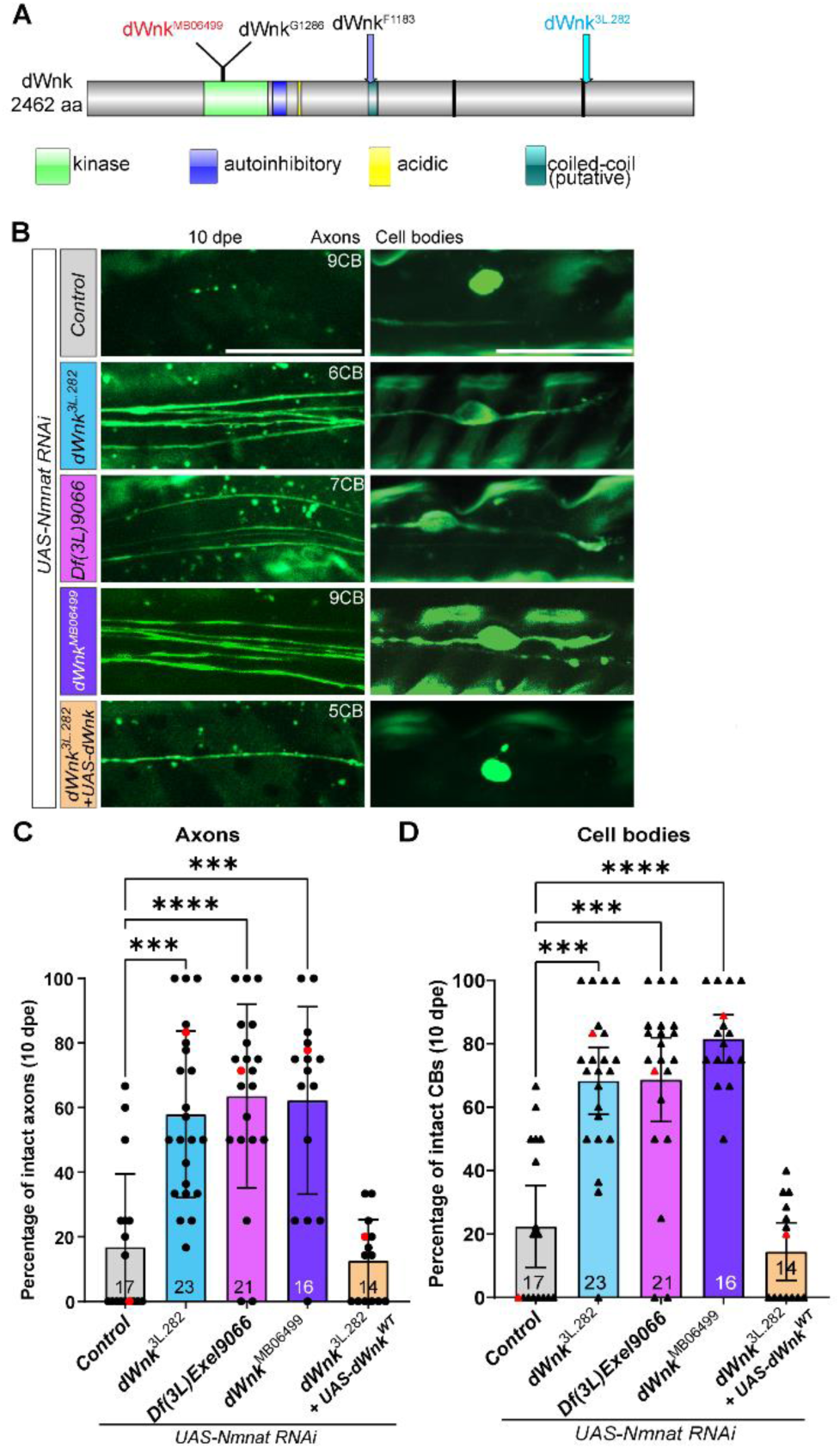
dWnk promotes neurodegeneration after dNmnat loss. **(A)** Schematic representation of the longest dWnk isoform (dWnk-PM, 2462 aa) and domains. Mutations used in this study are indicated. **(B)** Representative images of 10 days post-eclosion (dpe) wings showing axons and cell bodies. Genotypes are as indicated. Scale bar is 20 µm. **(C- D)** Quantifications of intact axons (C) and cell bodies (D) shown in B. Red dots and triangles correspond to the representative images in B. The results are plotted as mean ± 95% confidence interval (CI) and analyzed using Kruskal-Wallis tests with post-hoc Dunn’s multiple comparison test. All comparisons are listed in the Supplementary table 1.

We screened 4,917 mutant lines on the left arm of chromosome 3 in MARCM clones for suppression of dNmnat^RNAi^-induced neurodegeneration. Among several candidate mutations, we identified two lines, *3L.282* and *3L.1541*, which resulted in strong suppression of neurodegeneration, with 57.9% and 63.9% of neurons remaining intact, respectively, compared to 18.6% in controls (Fig. 1B-D and S1F). Cell bodies in these lines were slightly more protected than axons (Fig. 1D), suggesting that axonal degeneration might be more sensitive to dNmnat loss than cell death. Lines *3L.282* and *3L.1541* were lethal when crossed together, and both were lethal when crossed to either of two molecularly defined genomic deletions, *Df(3L)ED4978*^21^ and *Df(3L)Exel9066.* Both deficiencies delete the *Drosophila with-no-lysine kinase* (*dWnk*) locus along with additional genes (Fig. S1B, S1C). Whole genome sequencing of the *3L.282* line revealed a C-to-T mutation in *dWnk* that alters a splice donor site in intron nine, affecting all four *dWnk* transcripts (Fig. S1B). Sequencing of *3L.1541* did not reveal any mutations in the *dWnk* coding region, suggesting this mutation could affect a regulatory region of *dWnk*. To confirm that loss of dWnk caused the suppression of neurodegeneration after *dNmnat* knockdown in these mutant lines, we assayed two available molecularly defined null alleles of *dWnk* in MARCM clones, *Df(3L)Exel9066* and *dWnk^MB06499^* ^22^ (Fig. 1A and S1B). We found they both exhibited similar preservation of axons and cell bodies when compared to mutants *3L.282* and *3L.1541* (Fig. 1B- D). In addition, reintroducing wild-type *dWnk* (*UAS-dWnk^WT^*) into MARCM clones in these backgrounds restored neurodegeneration upon dNmnat depletion (Fig. 1B-D, and S1F). These data indicate that *3L.282* and *3L.1541* are loss-of-function mutations in *dWnk*, and that dWnk is required to drive neurodegeneration after dNmnat loss.

A previous study reported that other loss-of-function alleles, *dWnk^F^*^118^*^3^* and *dWnk^G1286^* (Fig. S1B, C), resulted in opposite effects, with highly branched pSC and pDC mechanosensory neurons showing defects in axon pathfinding and spontaneous neurodegeneration that could be suppressed by mutation of *dSarm* or *axed*^23^. Surprisingly, the above *dWnk* alleles did not protect against neurodegeneration induced by *dNmnat* depletion in our hands (Fig. S1G), suggesting they might be weaker, partial loss-of-function alleles. To investigate whether knocking out dWnk led to spontaneous degeneration in either the glutamatergic L1 wing sensory neurons or an orthogonal set of neurons, the cholinergic OR22a^+^ olfactory receptor neurons, we analyzed homozygous neuronal clones of various *dWnk* alleles in these cell types, including *dWnk^F1183^* and *dWnk^G1286^*. We found both neuron types remained morphologically intact late into adulthood (Fig. S1H). This discrepancy in functional roles for dWnk in cell maintenance versus activation of dSarm-sensitive degenerative events could point to a cell-specific role for dWnk in the development and maintenance of pSC and pDC (see also Discussion).

### dWnk kinase activity is required for pro-degenerative signaling

dWnk, like mammalian WNKs (WNK1-4), is a kinase that acts as a sensor for ions such as chloride and potassium and changes in osmotic pressure (Fig. 1A)^24,25^. We sought to determine which of these functions might regulate dWnk signaling in the context of neurodegeneration after dNmnat depletion. To determine whether the kinase and Cl^-^ binding activities of dWnk are required for its pro-degenerative role after dNmnat loss, we used a number of *UAS*-driven transgenes to overexpress modified forms of dWnk: an S632A mutation that disrupts auto- activation of dWnk via phosphorylation^26,27^, a D618A mutation that generates a dominant-negative dWnk lacking kinase activity^22,28,29^, and an L619F mutation that reduces the Cl^-^ sensitivity of dWnk, rendering it constitutively active^30^ (Fig. 2A). We first expressed these in wild type clones and found that constitutive expression of *dWnk^S632A^* or *dWnk^L619F^* did not affect neuron survival while expression of *dWnk^D618A^* did result in a slight reduction in cell survival (Fig. S2A). We next generated dNmnat^RNAi^-expressing MARCM clones and expressed each transgene and assayed for suppression of neurodegeneration. We found that expression of *dWnk^S632A^*did not suppress neurodegeneration after dNmnat depletion (Fig. 2B and S2B). In contrast, expression of the dominant-negative *dWnk^D618A^* molecule strongly suppressed neurodegeneration in the dNmnat^RNAi^ background, providing additional evidence for a pro-degenerative role for dWnk after dNmnat depletion (Fig. 2B and S2B). Finally, expression of the constitutively activated, Cl^-^- insensitive *dWnk^L619F^* molecule neither suppressed nor accelerated neurodegeneration after dNmnat loss (Fig. 2B and S2B). These observations suggest that while dWnk is essential for promoting neurodegeneration triggered by dNmnat depletion, overactivation of the pathway does not accelerate neurodegeneration.

**Figure 2.**
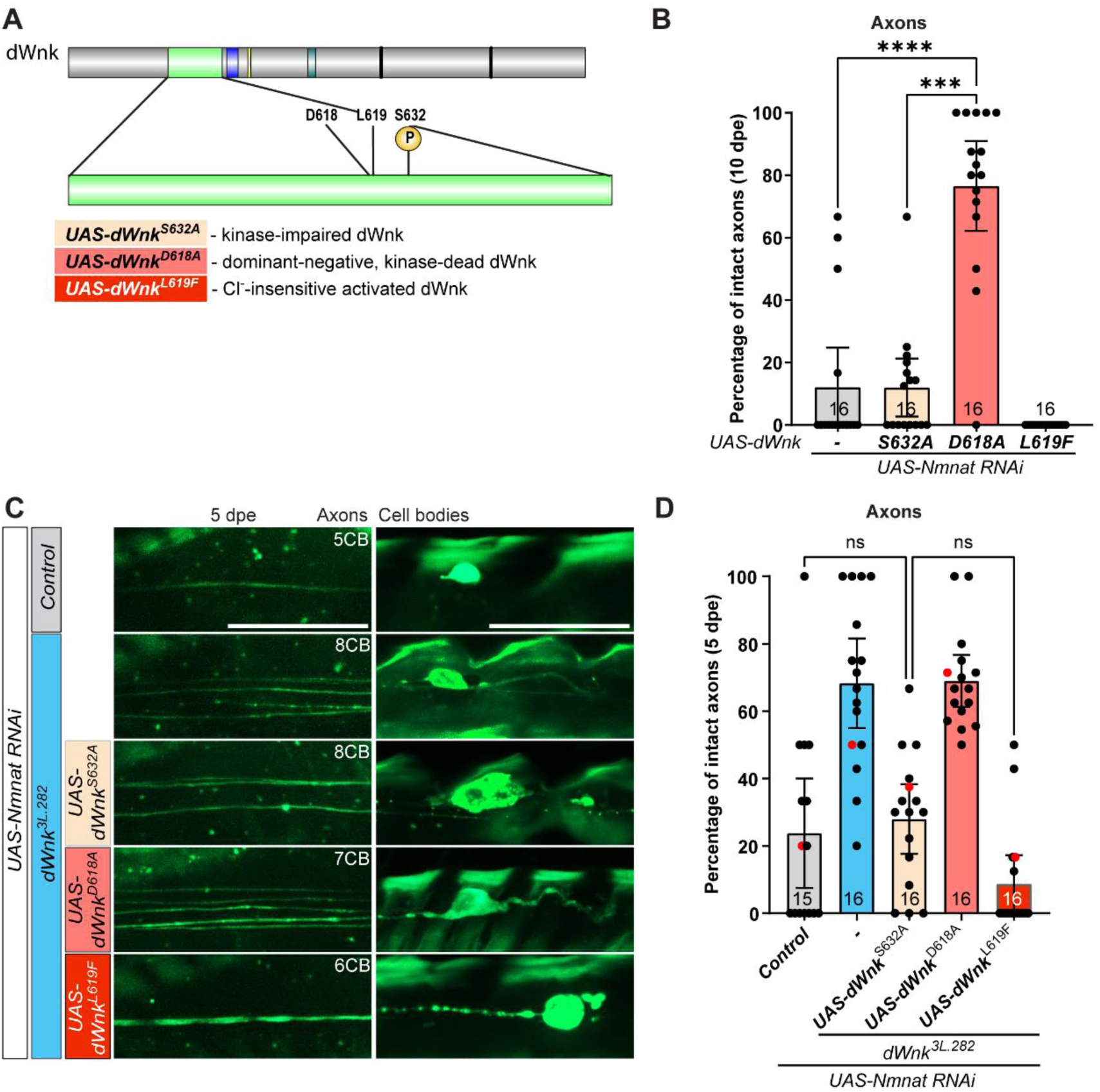
dWnk kinase activity is essential for driving neurodegeneration. **(A)** Schematic representation of the kinase domain of dWnk, with the residues important for dWnk kinase activity: D618 – ATP binding, L619 – chloride sensing, and S632 – first auto- phosphorylation site. **(B)** dWnk^D618A^ is a dominant-negative form of dWnk. Quantifications show the neuroprotection (10 dpe), genotypes as indicated. The results are plotted as mean ± 95% CI and analyzed using Kruskal-Wallis tests with post-hoc Dunn’s multiple comparison test. All comparisons are listed in the Supplementary table 1. **(C)** Representative images of 5 dpe wings, genotypes as indicated. Scale bar is 20 µm. **(D)** Quantifications of the experiments shown in C. The red dots correspond to the representative images in C. The results are plotted as mean ± 95% CI and analyzed using Kruskal-Wallis tests with post-hoc Dunn’s multiple comparison test. All comparisons are listed in the Supplementary table 1.

To determine whether each of these dWnk variants could rescue the blockade of neurodegeneration by *dWnk^3L.282^* after dNmnat depletion. We found that expression of *dWnk^S632A^*or *dWnk^L619F^* were sufficient to rescue *dWnk^3L.282^*mutant phenotypes, while expression of the dominant negative *dWnk^D618A^*molecule was not (Fig. 2C, D and S2C-E). We observed similar phenotypes when we examined the survival of neuronal cell bodies (Fig. S2C, E), and in the *Df(3L)Exel9066* and *dWnk^MB06499^* backgrounds with *dWnk^S632A^* (Fig. S2F, G). Together, these data argue the auto-activation by phosphorylation and Cl^-^ sensing functions of dWnk are not required for pro-degenerative signaling while dWnk kinase activity is essential, and that activation of dWnk signaling is sufficient to drive neurodegeneration after dNmnat depletion but not in healthy control neurons.

### Frayed acts downstream of dWnk to promote neurodegeneration

Mammalian WNK kinases signal through downstream kinases SPAK/OSR1, which in *Drosophila* are encoded by a single ortholog gene, *frayed* (*fray*)^31^ (Fig. S3B). WNKs phosphorylate SPAK/OSR1/Fray at specific sites that lead to the activation of downstream targets including cation-chloride co-transporters^28,29,32–35^ (Fig. 3B and S3B). We therefore sought to determine whether dWnk signals through Fray to drive neurodegeneration after dNmnat loss. For loss-of-function studies, we used the P-element insertion allele of *fray*, *fray^07551^* (Fig. S3A)^36,37^. When we crossed *fray^07551^* into a dNmnat^RNAI^ background, we observed strong neuroprotection, with ∼60% of neurons surviving in homozygous mutant MARCM clones (Fig. 3A, D, E). We confirmed this phenotype by generating two additional alleles of *fray*: a genome engineered deletion (*fray^Δ^*), and knock-in allele that eliminated the catalytic lysine at position 67 (*fray^K67M^*)^28^. We found that, while the *fray^Δ^* allele suppressed neurodegeneration activated by dNmnat depletion in MARCM clones, the kinase-impaired allele *fray^K67M^*exhibited reduced neuroprotection compared to the null allele (Fig. 3C), supporting a role for Fray and its kinase activity in pro-degenerative signaling.

**Figure 3.**
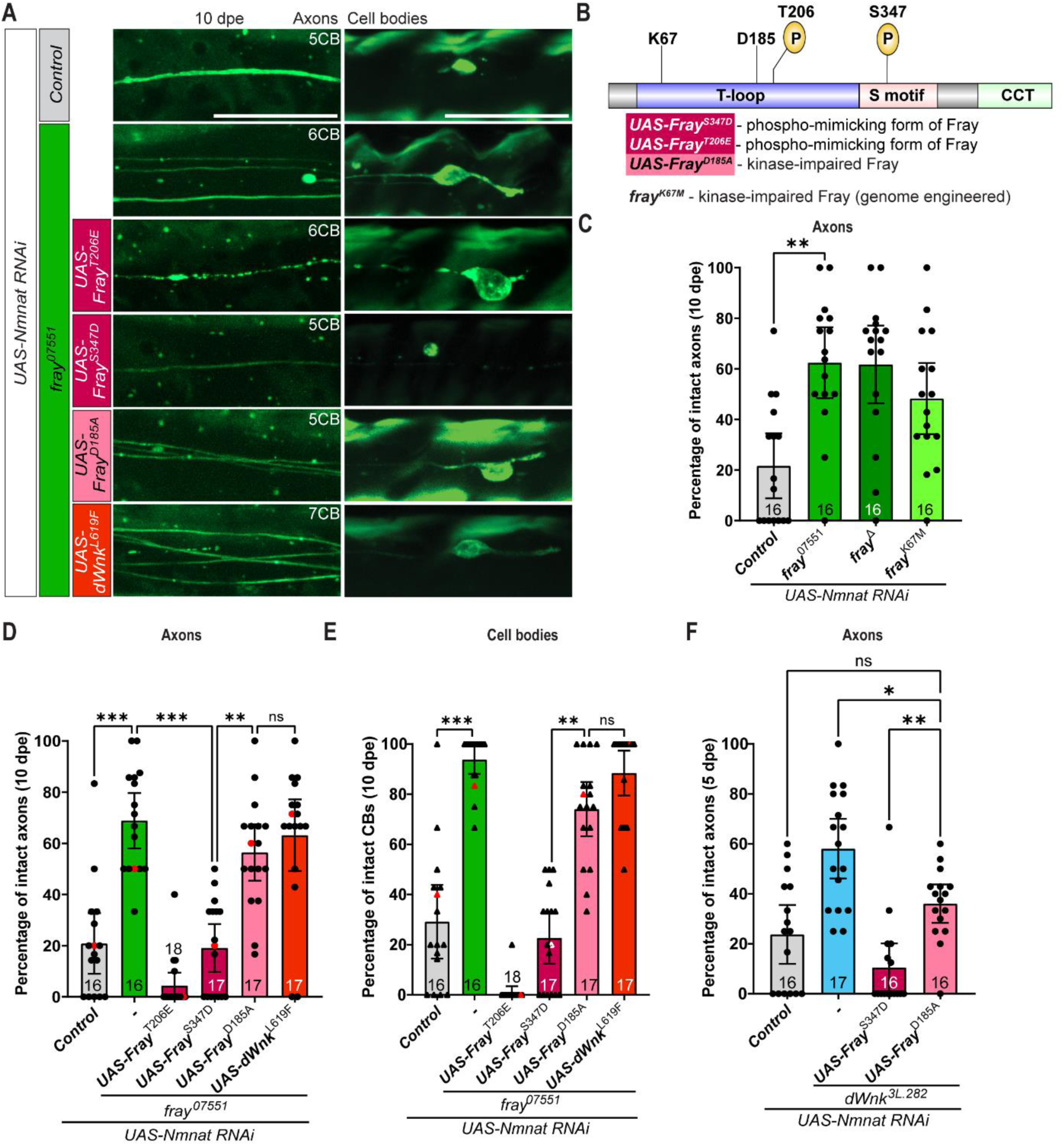
Fray kinase promotes neurodegeneration downstream of dWnk. **(A)** Representative images of 10 dpe wings showing axons and cell bodies, genotypes as indicated. Scale bar is 20 µm. **(B)** Schematic representation of Fray, indicating the residues important for Fray kinase activity: K67 – catalytic lysine, D185 – ATP binding, and T206 & S347 – dWnk phosphorylation sites. **(C)** Quantification for LOF alleles of *fray*, genotypes as indicated. The results are plotted as mean ± 95% CI and analyzed using Kruskal-Wallis tests with post-hoc Dunn’s multiple comparison test. All comparisons are listed in the Supplementary table 1. **(D-E)** Quantifications of intact axons (D) and cell bodies (E) shown in panel A. The red dots and triangles correspond to the representative images in A. The results are plotted as mean ± 95% CI and analyzed using Kruskal-Wallis tests with post-hoc Dunn’s multiple comparison test. All comparisons are listed in the Supplementary table 1. **(F)** Assay of the phospho-mimicking form of Fray, Fray^S347D^, kinase-impaired form of Fray, Fray^D185A^ (5 dpe) in *dWnk^3L.282^*mutants (5 dpe), genotypes as indicated. The results are plotted as mean ± 95% CI and analyzed using Kruskal- Wallis tests with post-hoc Dunn’s multiple comparison test. All comparisons are listed in the Supplementary table 1.

Fray has two key dWnk phosphorylation sites that drive activation, S347 and T206. We used the phospho-mimicking Fray^S347D^ and Fray^T206E^ versions of these, along with the kinase impaired Fray^D185A^, to explore how Fray signaling promoted neurodegeneration. We expressed each of these in *fray^07551^* MARCM clones expressing *dNmnat^RNAi^*and found that both Fray^S347D^ and Fray^T206E^ promoted rapid neurodegeneration while expression of the kinase-impaired Fray^D185A^ did not (Fig. 3A, D, E). Expression of each of these constructs in wild type MARCM clones in neurons did not result in neurodegeneration (Fig. S3H), suggesting that activation of Fray alone is insufficient to induce neurodegeneration and can only promote neurodegeneration after dNmant loss.

To determine whether Fray functions downstream of dWnk during neurodegeneration, we expressed an activated form of Fray (Fray^S347D^) in *dWnk* loss-of-function mutant backgrounds. As expected, we found Fray^S347D^ was sufficient to overcome the protective effects of *dWnk^3L.282^* and *dWnk^MB06499^*, while the kinase impaired Fray^D185A^ was not (Fig. 3F and S3C-G). In contrast, expression of the activated form of dWnk (dWnk^L619F^)^30^ in a *fray^07551^*background did not diminish the neuroprotection afforded by the *fray* loss-of-function mutant (Fig. 3A, D, E), indicating that Fray is required to drive dWnk-dependent neurodegeneration. In all cases, we observed similar phenotypes in axons and cell bodies in MARCM clones (Fig. 3D-F and S3C-G), suggesting that axons and cell bodies use dWnk and Fray signaling similarly to drive axon degeneration and cell death. In summary, these data provide strong support for a model whereby dWnk activation signals downstream to Fray to drive neurodegeneration induced by dNmnat loss.

### dWnk acts in parallel to dSarm to drive neurodegeneration

Our previous work demonstrated Axundead (Axed) functions downstream of dSarm in Wallerian degeneration^17^. Strikingly, loss of *axed* confers near-complete protection across various neurodegenerative paradigms, while *dSarm* nulls fail to protect in some neurodegenerative backgrounds, in particular neurodegeneration induced by *dNmnat* null mutations^17^. This observation strongly suggests that there are additional neurodegenerative pathways that are activated in parallel to dSarm and signal through Axed to bring on neuronal demise. To explore the relationship between dSarm, Axed, and the dWnk/Fray pathway, we utilized the temperature-sensitive *Gal4/UAS* expression system to induce partial knockdown of *dNmnat*. We conducted experiments at 29°C to increase *UAS-dNmnat^RNAi^* expression, resulting in more rapid neurodegeneration than at 25°C (our standard condition). In this context, 95% of *UAS- dNmnat^RNAi^*-expressing wild type MARCM clones degenerated by 8 dpe (Fig. 4A-C) compared to approximately 70% by 10 dpe at 25°C (Fig. S4A, B). Similarly, while at 25°C we observed 70% protection by a *dSarm* null allele (*dSarm^896^*)^9^ (Fig. S4A, B), at 29°C protection decreased to 46- 50% (Fig. 4A-C) and *dWnk* null mutant neuroprotection decreased from 57-60% at 25°C (Fig. S4A, B) to 31-38% at 29°C (Fig. 4A-C). Interestingly, when we then assayed *dSarm, dWnk* double mutants (*dSarm^896^*, *dWnk^3L.282^*), we found an additivity of protection, with 85% *dSarm^896^*, *dWnk^3L.282^* MARCM clones remaining intact (Fig. 4A-C), levels equivalent to *axed* null allele (*axed^3L.11^*)^17^ (Fig. 4A-C) alone. We found similar results using another double mutant combination, *dSarm^896^, dWnk^MB06499^*(Fig. S4C, D). Given these are null alleles, the additive nature of this phenotype indicates dWnk and dSarm act in parallel pathways to drive neurodegeneration in the context of dNmnat depletion.

**Figure 4.**
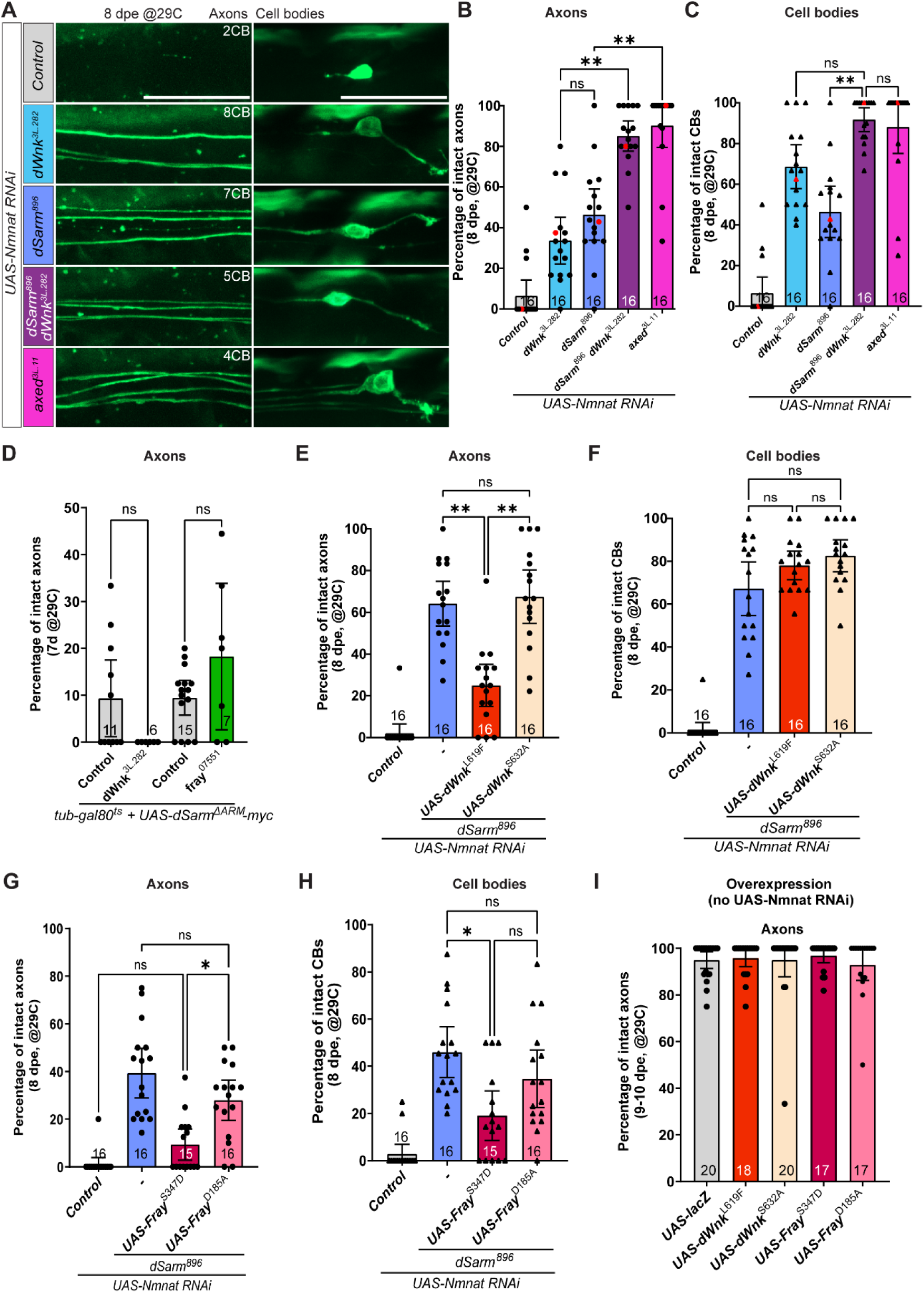
dWnk acts in parallel to dSarm to drive neurodegeneration after dNmnat loss. **(A)** Representative images of 8 dpe wings after *dNmnat KD*-induced neurodegeneration (29°C), genotypes as indicated. **(B-C)** Quantifications for (A): axonal protection is shown in panel B, and cell body protection in panel C. Red dots (B) and triangles (C) correspond to representative images in panel A. The results are plotted as mean ± 95% CI and analyzed using Kruskal-Wallis tests with post-hoc Dunn’s multiple comparison test. All comparisons are listed in the Supplementary table 1. **(D)** Assays of *dWnk* or *fray* mutants after dSarm^ΔARM^ expression. The results are plotted as mean ± 95% CI and analyzed using Kruskal-Wallis tests with post-hoc Dunn’s multiple comparison test. All comparisons are listed in the Supplementary table 1. **(E-F)** Assays of Cl⁻-insensitive dWnk (dWnk^L619F^) and auto-activation impaired dWnk (dWnk^S632A^) *dSarm^896^*backgrounds, genotypes as indicated. Axons versus cell bodies indicated. The results are plotted as mean ± 95% CI and analyzed using Kruskal-Wallis tests with post-hoc Dunn’s multiple comparison test. All comparisons are listed in the Supplementary table 1. **(G-H)** Expression of the phospho-mimicking form of Fray (Fray^S347D^), and kinase-impaired form (Fray^D185A^), in *dSarm^896^* neurons, genotypes and manipulations as indicated. The results are plotted as mean ± 95% CI and analyzed using Kruskal-Wallis tests with post-hoc Dunn’s multiple comparison test. All comparisons are listed in the Supplementary table 1. **(I)** Overexpression of dWnk^L619F^, dWnk^S632A^, Fray^S347D^ and Fray^D185A^ without *dNmnat KD*. Results are plotted as mean ± 95% CI. Data analysis is provided in the Supplementary table 1.

Can activation of dSarm or dWnk/Fray signaling bypass protection conferred by inhibition of the other pathway? To test this possibility, we expressed a gain-of-function form of dSarm that lacks the auto-inhibitory ARM domain (*dSarm^ΔARM^*)^17,38^. This resulted in rapid neurodegeneration that was unaffected by either *dWnk* or *fray* null mutations (Fig. 4D). In contrast, while expression of the activated version of dWnk (dWnk^L619F^) or Fray (Fray^S347D^) did not trigger spontaneous neurodegeneration at 29°C on their own in control MARCM clones (Fig. 4I), expression of either in *dNmnat^RNAi^* neurons significantly reduced the protection offered by *dSarm^896^* (Fig. 4E-H). However, expressing kinase-impaired forms of either dWnk or Fray could not override protection provided by *dSarm* nulls (Fig. 4E-H). Together, these data indicate dWnk and Fray act in parallel to dSarm, with activation of either pathway being sufficient to overcome the other in the context of neurodegeneration after dNmnat loss (Fig. 5B).

**Figure 5.**
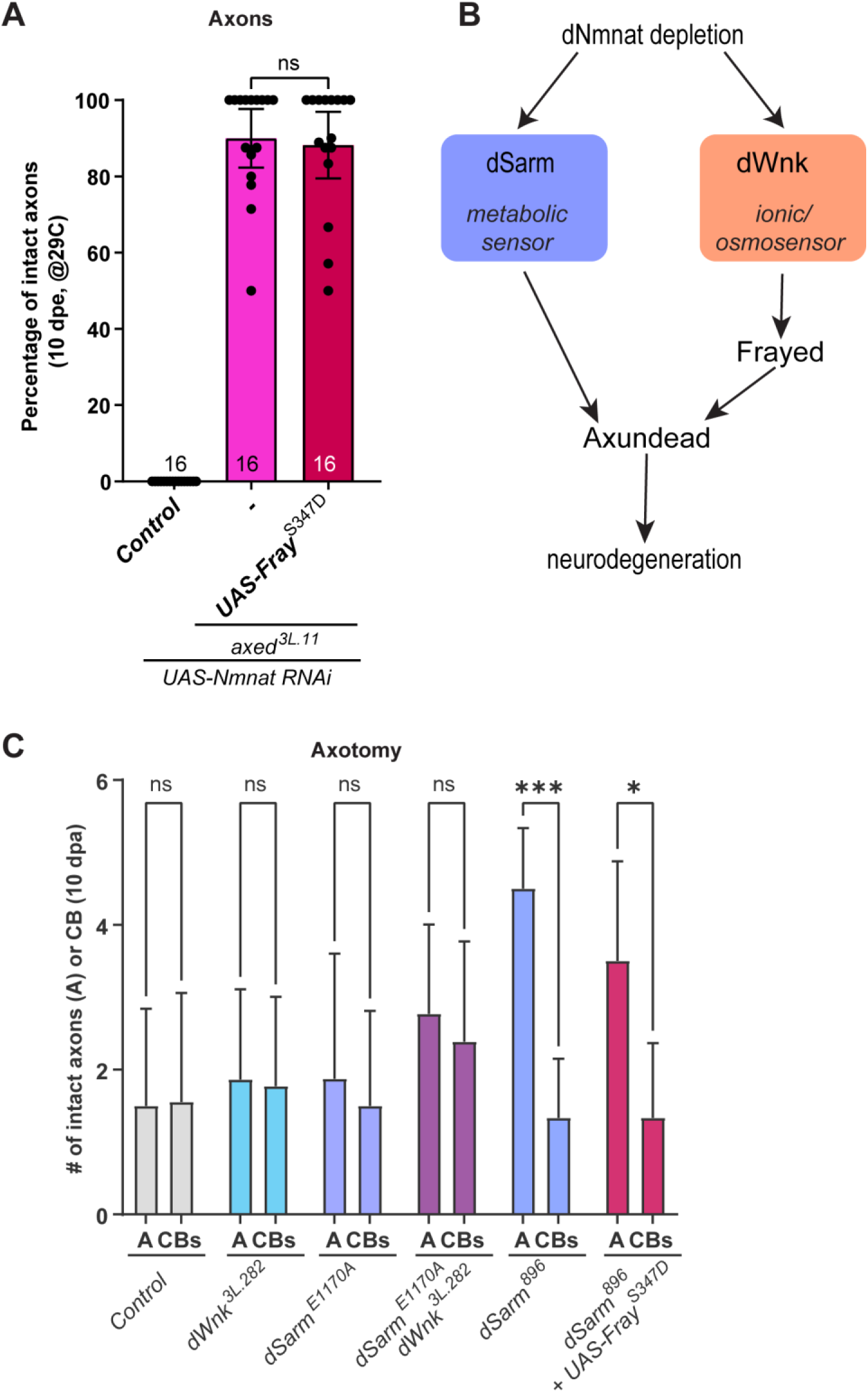
dWnk/Fray and dSarm converge on Axed, but dWnk/Fray loss does not block Wallerian degeneration. **(A)** Controls and expression of the phospho-mimicking form of Fray, Fray^S347D^, in *axed^3L.11^*mutants. The results are plotted as mean ± 95% CI and analyzed using Kruskal-Wallis tests with post-hoc Dunn’s multiple comparison test. All comparisons are listed in the Supplementary table **(B)** Model for parallel activation of dWnk/Fray and dSarm, upstream of Axed. **(C)** Axonal protection in controls, *dWnk*, *dSarm*, and with the phospho-mimicking form of Fray, Fray^S347D^ (axotomy). The results are plotted as mean ± SD and analyzed using 2way ANOVA with Šídák’s multiple comparisons test. *** - p-value 0.0001 to 0.001, * - p-value 0.01 to 0.05, ns - p-value ≥ 0.05.

### dWnk/Fray and dSarm converge on Axed to drive neurodegeneration

We next sought to identify the effectors downstream of Fray that promote neurodegeneration. We assessed roles for molecules downstream of dWnk/Fray in other contexts, such as the Na^+^/K^+^/Cl^-^ co-transporters (*NKCC1*/2 in mammals, *Ncc69* and *NKCC* in *Drosophila*) in ion balance and cell volume regulation^25^ and inward-rectifier potassium channels (Kir in mammals, Irk in *Drosophila*) that regulate circadian behaviors^35^ (Fig. S3B). Loss-of-function alleles of *Ncc69, Irk1*, *Irk2* and *Irk3* did not protect neurons against dNmnat depletion-induced neurodegeneration (Fig. S5A, B). Fray-mediated dWnk signaling may therefore be executed by different downstream signaling molecules, although we cannot exclude the possibility that the above molecules act in a genetically redundant fashion to drive axon destruction, which would have been missed in our assays conducted with single loss-of-function mutants.

We next asked whether dWnk and Fray promote neurodegeneration via Axed. To this end, we expressed an activated version of Fray (Fray^S347D^) in an *axed* null mutant background expressing dNmant^RNAi^. Interestingly, although expression of Fray^S347D^ was sufficient to overcome the protection afforded by *dSarm* nulls, activated Fray was not sufficient to overcome the strong protection afforded by *axed* mutants (Fig. 5A). This observation, coupled with the additivity of *dSarm* and *dWnk/Fray* null mutants, further supports a model whereby depletion of dNmnat drives dSarm and dWnk/Fray to act in parallel to drive neurodegeneration through Axed (Fig. 5B).

Although mutations in *dWnk* and *fray* can suppress neurodegeneration that is activated by dNmnat depletion, we found *dWnk* mutations did not block axon degeneration after axotomy (Fig. 5C). We explored the possibility that *dWnk* mutations might enhance *dSarm* mutant phenotypes after axotomy. In previous work, we generated a hypomorphic allele of *dSarm* predicted to lack NAD^+^ hydrolase activity, *dSarm^E1170A^*. Whereas *dSarm* null mutants protect severed axons for weeks after axotomy^9,17^, *dSarm^E1170A^* only protected severed axons for 5 days^39^. We crossed strong *dWnk* loss-of-function alleles into this background and severed axons but found that *dSarm^E1170A^, dWnk^3L.282^* MARCM clones displayed no increased protection after axotomy relative to *dSarm^E1170A^* alone at 10 days post-axotomy (dpa) (Fig. 5C). These findings imply that, despite dNmnat loss occurring in both contexts, dWnk loss is not sufficient to protect severed distal axons or enhance partial *dSarm* mutant phenotypes in the context of axotomy. Finally, given that expression of activated Fray^S347D^ can overcome *dSarm* null mutant neuroprotection against dNmnat depletion, we asked whether it would similarly bypass the protection afforded by *dSarm^896^* null mutations in distal severed axons after axotomy. Surprisingly, we found that Fray^S347D^ expression did not override the protection of distal severed axons in *dSarm^896^*null mutants even 10 dpa (Fig. 5C). We envision two possible explanations for these observations: dNmnat depletion and axotomy are not equivalent with respect to neurodegenerative signaling and dWnk/Fray are only required after dNmnat depletion; or the ability of dWnk/Fray signaling to promote neurodegeneration may require signaling through the neuronal cell body.

## Discussion

Our work demonstrates that dWnk and Fray act as executioners of neurodegeneration caused by dNmnat depletion, which is also a potent activator of the dSarm/SARM1 pro-degenerative pathway^6,17^. WNKs and Fray/SPAK/OSR1 are conserved regulators of neuronal development and function, and growing evidence points to a role for WNKs/SPAK/OSR1 in neurodegeneraiton. Ischemic stroke is characterized by extensive cell death in the ischemic core, which can activate the neuronal WNK pathway in the ischemic penumbra, perhaps in response to osmotic stress^40^. SARM1 has also recently been shown to promote secondary thalamic neurodegeneration after cerebral infarction^41^. Interestingly, deletion of *WNK3* or *SPAK* reduces tissue damage, including neuronal loss, by inhibiting NKCC1 activation in mouse models of stroke^42,43^. In a primary cortical neuron ischemia/reperfusion model, NKCC1-dependent sodium influx triggers the reverse mode of the Na^+^/Ca^2+^ exchanger (NCX)^44^, raising intracellular Ca^2+^ levels, a well-known driver of neurodegeneration. How dWnk/Fray might signal to promote neurodegeneration remains an open question. To our surprise, elimination of one of the two *Drosophila* NKCCs (Ncc69) or any of the three Irk channels – the other family of potassium channels regulated by the WNK pathway – individually did not replicate the phenotypes observed with *dWnk* or *fray* loss. However, we cannot exclude the possibility that these molecules act in a functionally redundant fashion and that phenotypes might be revealed in double or triple loss-of-function mutants.

Mutations in WNKs are lalso inked to neurodevelopmental disorders characterized by disrupted GABAergic signaling, including epilepsy, autism spectrum disorder, schizophrenia, and intellectual disability^45^. Hereditary sensory and autonomic neuropathy type II (HSANII), a rare neurodegenerative condition, is caused by mutations in the HSN2 exon of *WNK1*. These mutations result in a truncated WNK1 protein that retains the kinase domain but lacks the C-terminal RFxV/I motif, necessary for binding downstream targets^46,47^. In mouse models, removal of the HSN2 exon does not replicate the peripheral neuropathy seen in HSANII patients, suggesting that the presence of the truncated WNK1 protein may drive neuropathy^48^. The *Drosophila dWnk^F1183^* allele, which results in defects in axon outgrowth and maintenance^23,49^, also produces a protein with an intact kinase domain lacking the C-terminal Fray-binding motif, and mimics the mutations causing HSANII^46^ and disrupts visual system wiring^49^. Changes dWnk and WNK1 signaling caused by protein truncations could explain why we did not observe neurodevelopmental defects in clean dWnk null backgrounds, and why the *dWnk^F11^*^83^ allele failed to suppress neurodegeneration in our hands. It is also clear that dWnk signaling during development and in the mature nervous system are distinct: while Fray is required for all pro-degenerative activity of dWnk after dNmnat loss, neurodevelopmental defects associated with dWnk signaling did not require Fray^23^. How dWnk could promote neurodevelopmental or neurodegenerative phenotypes in the absence of Fray remains an open question.

Our data indicate dWnk/Fray act as a separate branch of the dSarm/Axed pro-degenerative signaling pathway. We propose a model whereby after dNmnat depletion, dWnk signals through Fray to trigger neurodegeneration in parallel to dSarm/SARM1, as a separate branch of the axon degeneration pathway, and that dWnk/Fray and dSarm signaling converge on Axed to promote neuronal destruction. Curiously, unlike dSarm/SARM1, activation of the dWnk/Fray signaling pathway is only capable of triggering neurodegeneration after dNmnat has been depleted, and not in control axons. How might this be regulated? The dWnk/WNK family of kinases are unique in their ability to directly sense changes in chloride and potassium ions or osmotic pressure^25^. The chloride binding site in the dWnk/WNK kinase domain is well-defined, with chloride binding directly inhibiting kinase activity^30^, but we found this domain is dispensable for dWnk pro-degenerative function after dNmnat depletion. However, intriguingly, the kinase domain from dWnk and mammalian WNK3 and WNK4 are also directly inhibited by high levels of K^+^ in the 80-180 mM range (i.e. in the range of physiological intracellular K^+^ in neurons)^50^, although the precise mechanism by which K^+^ inhibits WNK kinase activity is not defined. Given that loss of dNmnat/NMNAT2 (or axon damage) leads to a catastrophic decrease in NAD^+^ and ATP, we propose dWnk is normally inhibited by high physiological levels of intracellular K^+^ but is activated after failure of the Na^+^/K^+^ ATPase in axons after NAD+ and ATP depletion. In this way, dWnk/WNKs would serve as ionic sensors in axons during physiological stress, with the potential to activate catastrophic neurodegeneration. This activity would occur in parallel to the NAD^+^/NMN metabolic sensing that is mediated by dSarm/SARM1. Neurons therefore seem capable of continuously sensing both ionic (dWnk/WNK) and metabolic (dSarm/SARM1) status, and when sufficiently dysfunctional, potently activate neurodegeneration through Axed and its downstream regulators. Finally, our data suggests that the neuroprotective effects of dSarm/SARM1 blockade in neurological disease may be enhanced by simultaneous inhibition of WNKs or SPAK/OSR1.

## Materials and Methods

### KEY RESOURCES TABLE

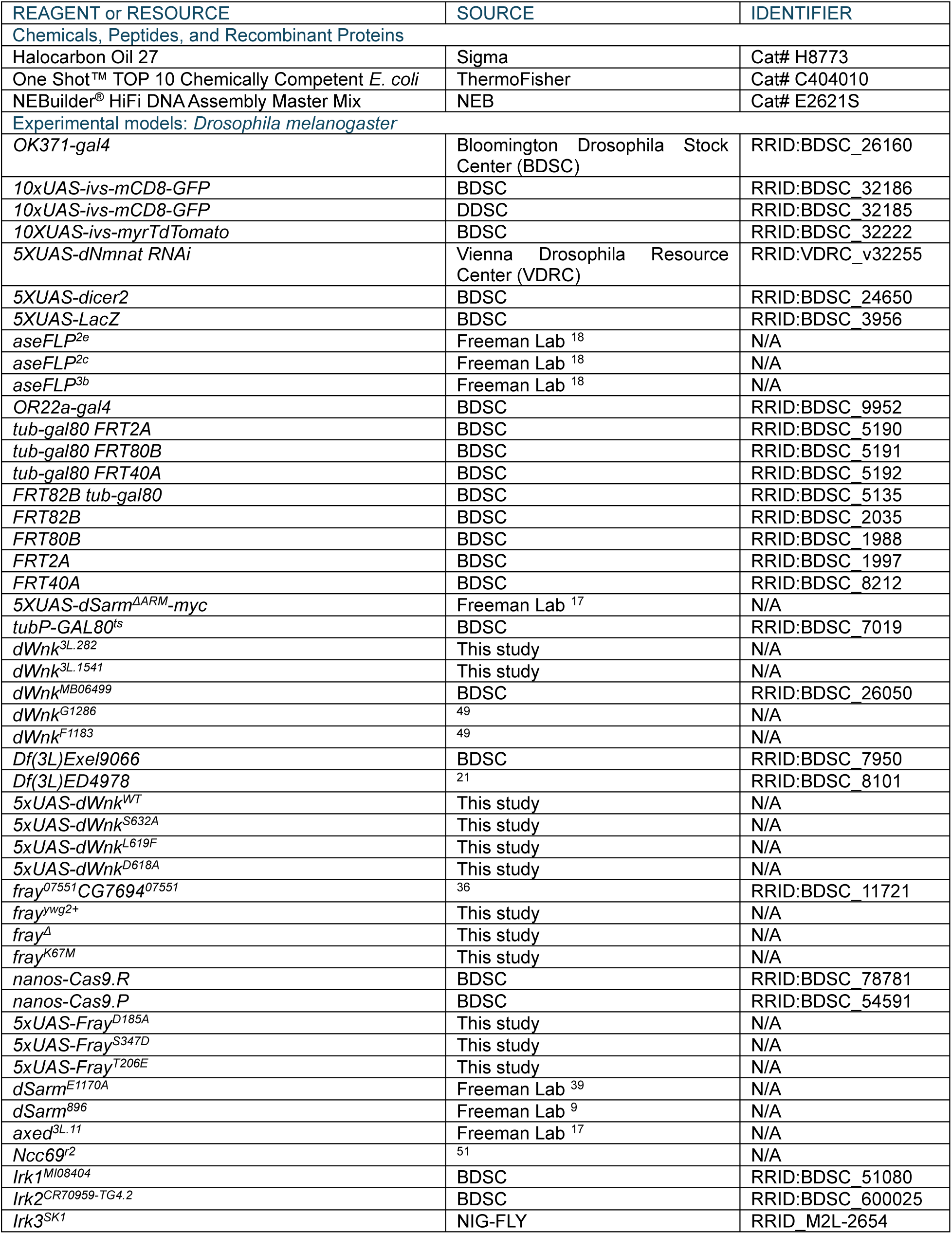
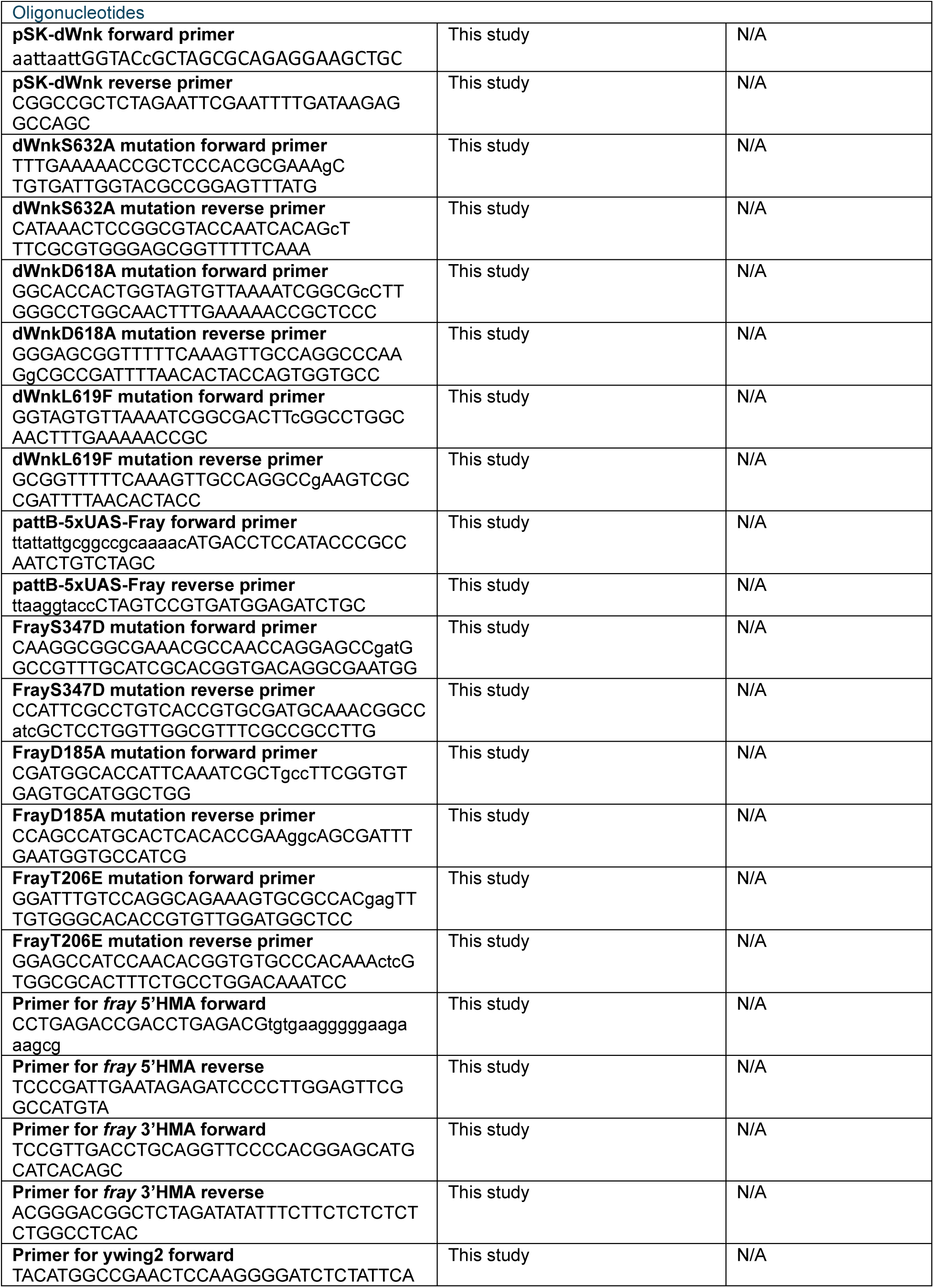
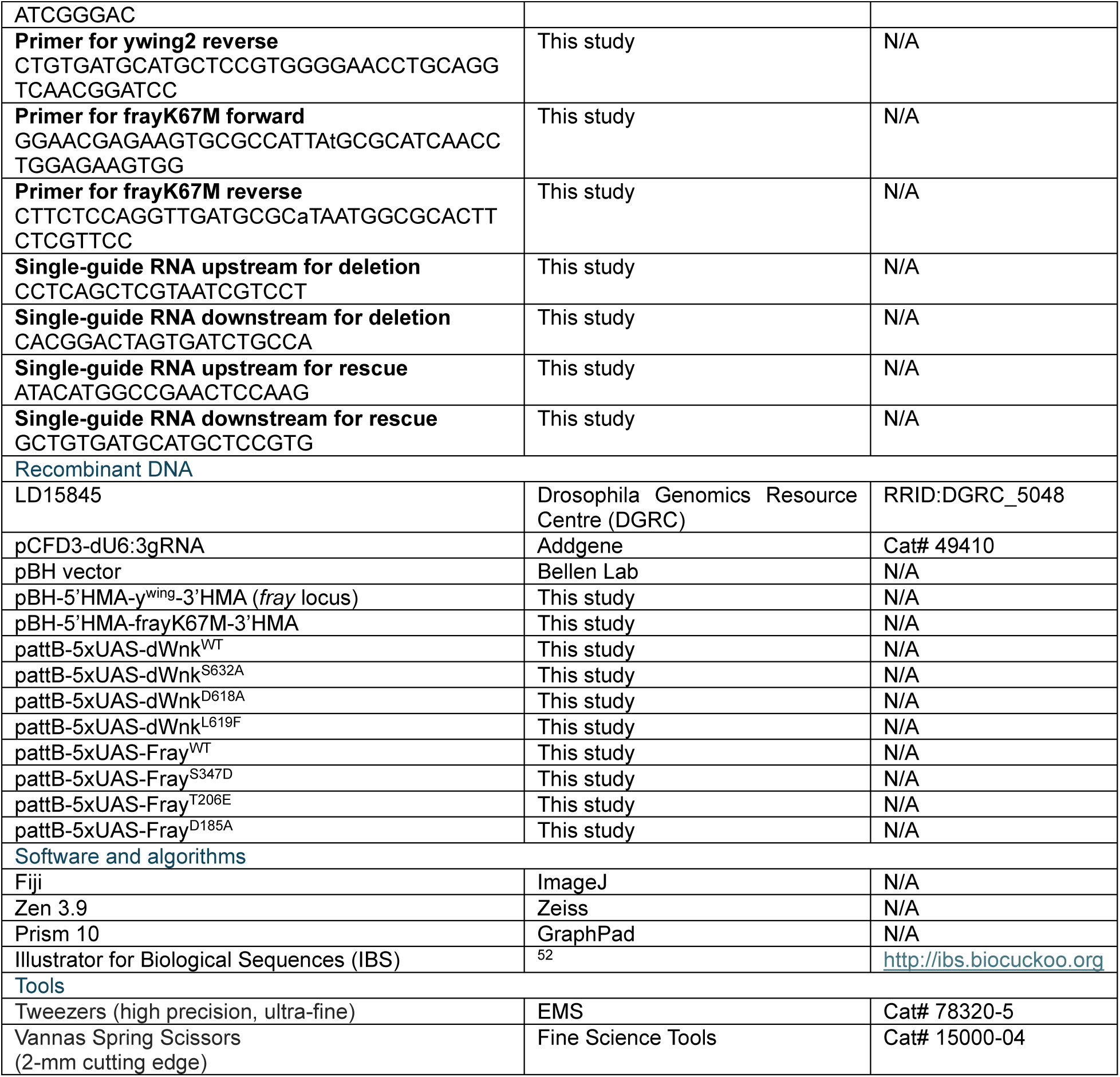

RESOURCE AVAILABILITY

## Leading contact

Further information and requests for resources and reagents should be directed to and will be fulfilled by the Lead Contact Marc R. Freeman.

## Materials availability

All unique reagents generated in this study are available from the Lead Contact without restriction.

## Data and code availability

This study did not generate any unique datasets or code.

## EXPERIMENTAL MODEL AND SUBJECT DETAILS

### Animals culture and temperature regimes

Flies (*D. melanogaster*) were maintained on standard cornmeal agar media supplemented with dry yeast at 25°C for the experiments presented in Figures 1-3 & 5C. For Figures 4 and 5A, flies were kept at 25°C for 3-4 days, after which the parents were transferred to fresh vials, and the progeny were raised at 29°C until imaging. For experiments involving the left arm of the second chromosome, *Irk3* mutant (Fig. S5A and B), crosses were maintained at 25°C, and eclosed animals were transferred to 29°C for 10 days until imaging. Equal numbers of males and females were included in each experiment. The complete list of fly lines used in this study is provided in the Key Resources Table, and the specific genotypes used for each figure are detailed in Supplementary Table 3.

## METHOD DETAILS

### Axotomy assays

Flies (5-6 days old) were anesthetized on a CO2 pad, and the L1 wing vein was cut at the midpoint, just above the posterior cross-vein, using Vannas Spring scissors. After the injury, the flies were returned to vials and incubated at 25°C. Wings were dissected for imaging 10 days post-axotomy. For the *dSarm^896^ + UAS-Fray^S347D^* experiment, flies were raised at 29°C for 8 days before the L1 wing vein was cut. After the injury, flies were returned to vials and incubated at 25°C. The wings were dissected for imaging 10 days post-axotomy.

### dSarm^ΔARM^ expression in adults

Virgin females of *5xUAS-dSarm^ΔARM^-myc*, *tub-gal80^ts^/CyO*; *tub-gal80 FRT2A* (or *FRT82B tub- gal80*)*/TM3, Sb*^1^ genotype were crossed to the appropriate MARCM males bearing *dWnk^3L.282^* or *fray^07551^* alleles, respectively. The crosses were incubated at 18°C (where Gal80^ts^ is active, thus halting Gal4-mediated expression). Eclosed progeny of the correct genotype were shifted to 29°C (where Gal80^ts^ is inactive) to induce Gal4-mediated expression of dSarm^ΔARM^. Complete neurodegeneration in control backgrounds was observed after 7 days at 29°C, which was selected as the experimental time point.

### 5xUAS-dWnkWT, 5xUAS-dWnk^S632A^, 5xUAS-dWnk^D618A^ and 5xUAS-dWnk^L619F^ cloning and transgenesis

The open reading frame of the longest dWnk transcript was synthesized by Genewiz and the *puc57-wnk* vector served as the starting template for all dWnk cloning. For targeted mutagenesis, base pairs 1259-2751 of the *dWnk* sequence, which contain all sites intended for mutation, were PCR-amplified while incorporating *KpnI* and *XbaI* sites at the 5’ and 3’ ends, respectively. The amplified fragment was then cloned into the *pSK* vector using *KpnI/XbaI* restriction sites. Specific mutations were introduced at desired sites within this region of *dWnk* using the QuikChange II-E Site-Directed Mutagenesis Kit. Primers used for this process are listed in the Key Resources Table. Mutagenized fragments were confirmed by sequencing, after which the targeted fragments were excised using *NheI* and *BstBI* restriction enzymes and cloned back into the *puc57-wnk* vector. To insert *dWnk* sequences into the *pattB-5xUAS* vector, the *dWnk* sequence was removed from *puc57-wnk* using *NotI* and *KpnI* restriction enzymes and then cloned into the *pattB-5xUAS* vector. The DNA was injected into *y^1^ w^67c23^ P{CaryP}attP18* (RRID:BDSC_32107) and *y1 w^67c23^; P{CaryP}attP154* flies by BestGene.

### 5xUAS-FrayWT, 5xUAS-Fray^D185A^, 5xUAS-Fray^S347D^ and 5xUAS-Fray^T206E^ cloning and transgenesis

To create Fray constructs, the *pBluescript (BS)-Fray* (*LD15845* cDNA clone, see Key Resources Table) was used as a template. To generate the *Fray^WT^* construct, Fray sequence was PCR-amplified from *pBS-Fray* with adding a *NotI* restriction site and a Kozak sequence to the 5’ end and a *KpnI* site to the 3’ end. The amplified *Fray* fragment was then cloned into the *NotI* and *KpnI* sites of the *pattB-5xUAS* vector. Specific mutations were introduced into *Fray* sequence using the QuikChange II-E Site-Directed Mutagenesis Kit. Primers used for this process are listed in the Key Resources Table. Once the mutation was confirmed, *Fray* sequence was excised from *pBS- Fray* using *NotI* and *KpnI* and cloned it into the *pattB-5xUAS* vector. The DNA was injected into *y^1^ w^67c23^ P{CaryP}attP18* (RRID:BDSC_32107) and *y1 w^67c23^; P{CaryP}attP154* flies by BestGene.

### Generation of *fray^K67M^* and *fray^Δ^* flies

The *fray^K67M^* and *fray^Δ^* fly lines were generated following a previously described protocol ^53^. A 1560 bp region containing the DNA sequence of *fray* exon 2 and 7 bp of the 3’UTR was selected for replacement with the *ywing2+* cassette. To construct the donor DNA (*pBH-5’HMA-p{ywing2+}- 3’HMA*), the homology arms (HMA) were PCR-amplified from the genomic DNA of *y^1^ sc* v^1^ sev^21^; P{nanos-Cas9.R}attP40/CyO* flies. Primers used for this process are listed in the Key Resources Table. The homology arms, *pBH* donor vector, and *p{y^wing2+^}* were combined using NEBuilder^®^ HiFi DNA Assembly Master Mix with the *BbsI* restriction enzyme. The final product was confirmed by sequencing.

For the generation of sgRNA, standard cloning protocols were followed using the *pCFD3- dU6:3gRNA* plasmid (https://www.crisprflydesign.org). Briefly, sense and antisense oligos containing the 20 bp guide target sequence were annealed and cloned into the *BbsI*-cut *pCFD3- dU6:3gRNA* plasmid. To generate the *fray^ywg2+^* flies, a mixture of 25 ng/ml sgRNA and 150 ng/ml donor DNA was injected into *y^1^ sc* v^1^ sev^21^; P{nanos-Cas9.R}attP40/CyO* flies by BestGene. The injected animals were then crossed to *yw* flies. Offspring were screened for yellow+ wings and crossed to the appropriate balancer flies. The entire homology arm sequence and about 200 bp of adjacent cassette sequences were verified by sequencing. Verified stocks were recombined to the *FRT82B* chromosome and tested for protection against *dNmnat KD*.

To replace the *y^wing2+^* cassette with the *frayK67M* sequence, donor DNA containing the *K67M* mutation (*pBH-5’HMA-frayK67M-3’HMA*) was generated (primers listed in the Key Resources Table). Briefly, primers containing the mutated nucleotides were used to amplify DNA fragments (*5pHMA-frayK67M* for the first half and *frayK67M-3pHMA* for the second half). NEBuilder^®^ HiFi DNA Assembly Master Mix was used to combine all DNA fragments with the *BbsI*- cut *pBH* vector. sgRNA was generated as described above, using different 20 bp guide target sequences (Key Resources Table). A mixture of 25 ng/ml sgRNA and 150 ng/ml donor DNA was injected into *y^1^ M{w^+mC^=nos-Cas9.P}ZH-2A w*;; FRT82B fray^ywg2+^/TM3,Sb* flies by BestGene. Progeny were screened for the loss of the yellow+ wing phenotype. These flies were balanced, and the entire replaced region was sequenced to confirm the absence of additional mutations. One of the screened stocks exhibited the complete removal of the *y^wing2+^* cassette without replacement by the *frayK67M* donor DNA, thus generating a null allele (*fray^Δ^*).

### Quantification and Statistical Analysis

Data acquisition and quantification were conducted in a non-blinded manner. Statistical analyses were performed using GraphPad Prism 10. Statistical details (including test definitions, number of wings analyzed, mean ± 95% CI, and p-values) are provided in the figure legends and Supplementary Tables 1 and 2. When meiotic recombination was used to generate a line, at least two independent recombinants were tested.

## ACKNOWLEDGEMENTS

We thank all the lab members of Freeman lab for the suggestions and discussions on this work and manuscript writing. We thank Prof. Aylin Rodan in the University of Utah for sharing ncc69^r1^ and ncc69^r2^ flies. We thank Prof. Dietmar Schmucker at the University of Bonn, Germany for sharing dWnk^F1183^ and dWnk^G1286^ flies and for discussions. We thank Dr. Li-Kroeger in Dr. Hugo Bellen lab in Baylor College of Medicine for sharing pBH vector. This work was supported by a generous gift from ALS Finding a Cure, and from NIH RO1 NS059991 (to MRF).

---

**Figure S1.**
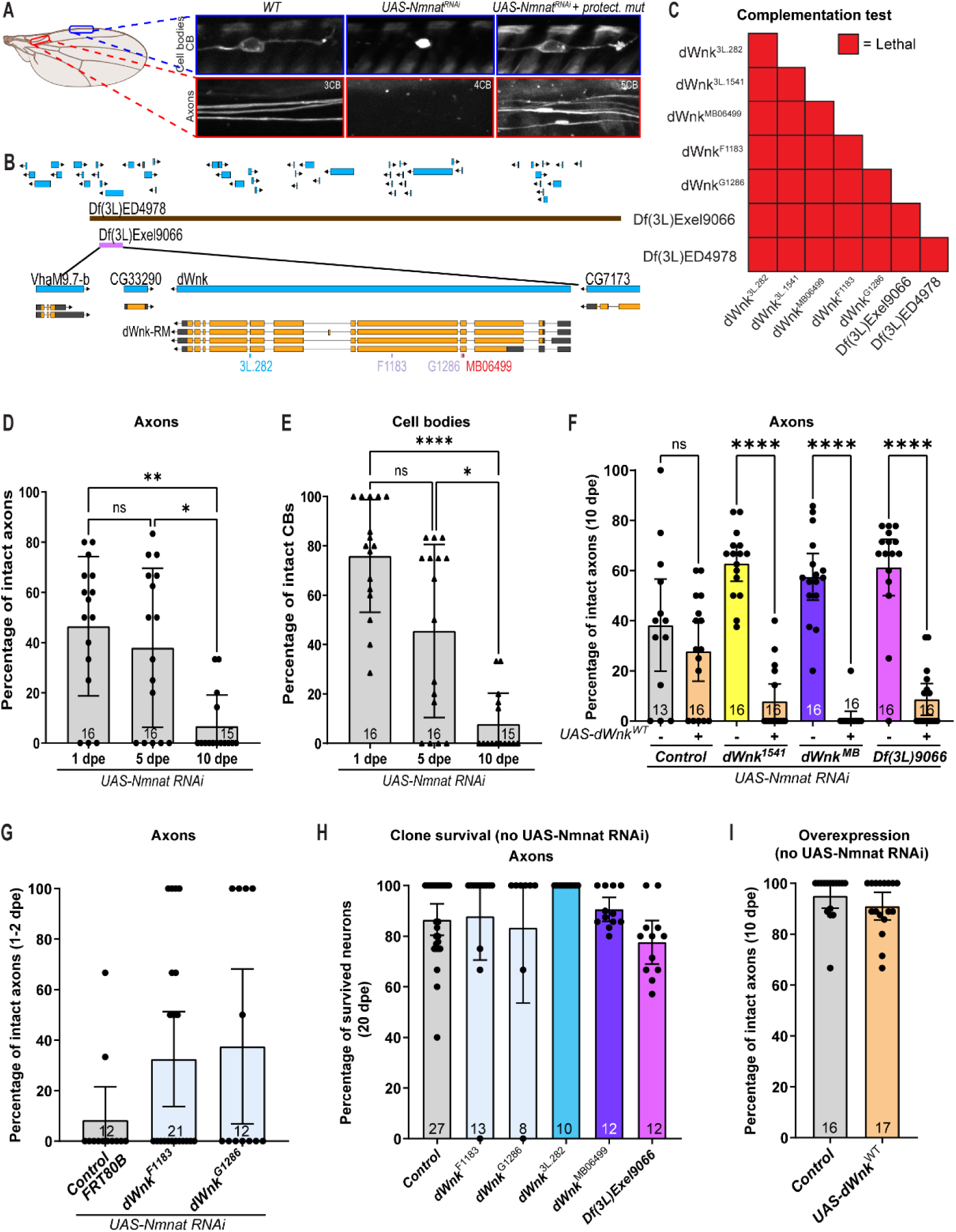
Related to Figure 1. **(A)** Schematics of the *dNmnat KD* screen: red insets represent images of distal axons, and blue insets represent images of neuronal cell bodies. dNmnat depletion drives neurodegeneration. Protective backgrounds offer protection to both axons and cell bodies. **(B)** Schematic representation of the *dWnk* genomic locus, showing the genomic deletions and *dWnk* alleles used in this study. **(C)** Complementation tests with LOF and null alleles of *dWnk* indicate that *dWnk^3L.282^*and *dWnk^3L.1541^* are LOF alleles. **(D-E)** Quantification of axonal (D) and cell bodies (E) survival following dNmnat depletion at 1, 5 and 10 dpe. The results are plotted as mean ± 95% CI and analyzed using Kruskal-Wallis tests with post-hoc Dunn’s multiple comparison test. All comparisons are listed in the Supplementary table 2. **(F)** Re-supply of dWnk^WT^ into *dWnk^3L.1541^*, *dWnk^MB06499^*, and *Df(3L)Exel9066* neurons restores their ability to degenerate following dNmnat depletion. The results are plotted as mean ± 95% CI and analyzed using Kruskal-Wallis tests with post-hoc Dunn’s multiple comparison test. All comparisons are listed in the Supplementary table **(G)** *dWnk^G1286^* and *dWnk^F1183^* alleles did not provide neuroprotection against dNmnat depletion. The results are plotted as mean ± 95% CI. Data analysis is provided in the Supplementary table **(H)** Neurons homozygous for *dWnk^G1286^*, *dWnk^F1183^*, *dWnk^3L.282^*, *dWnk^MB06499^*and *Df(3L)Exel9066* do not degenerate at 20 dpe. The results are plotted as mean ± 95% CI. Data analysis is provided in the Supplementary table 2. **(I)** Overexpression of dWnk^WT^ without *dNmnat KD* does not cause neurodegeneration. The results are plotted as mean ± 95% CI. Data analysis is provided in the Supplementary table 2.

**Figure S2.**
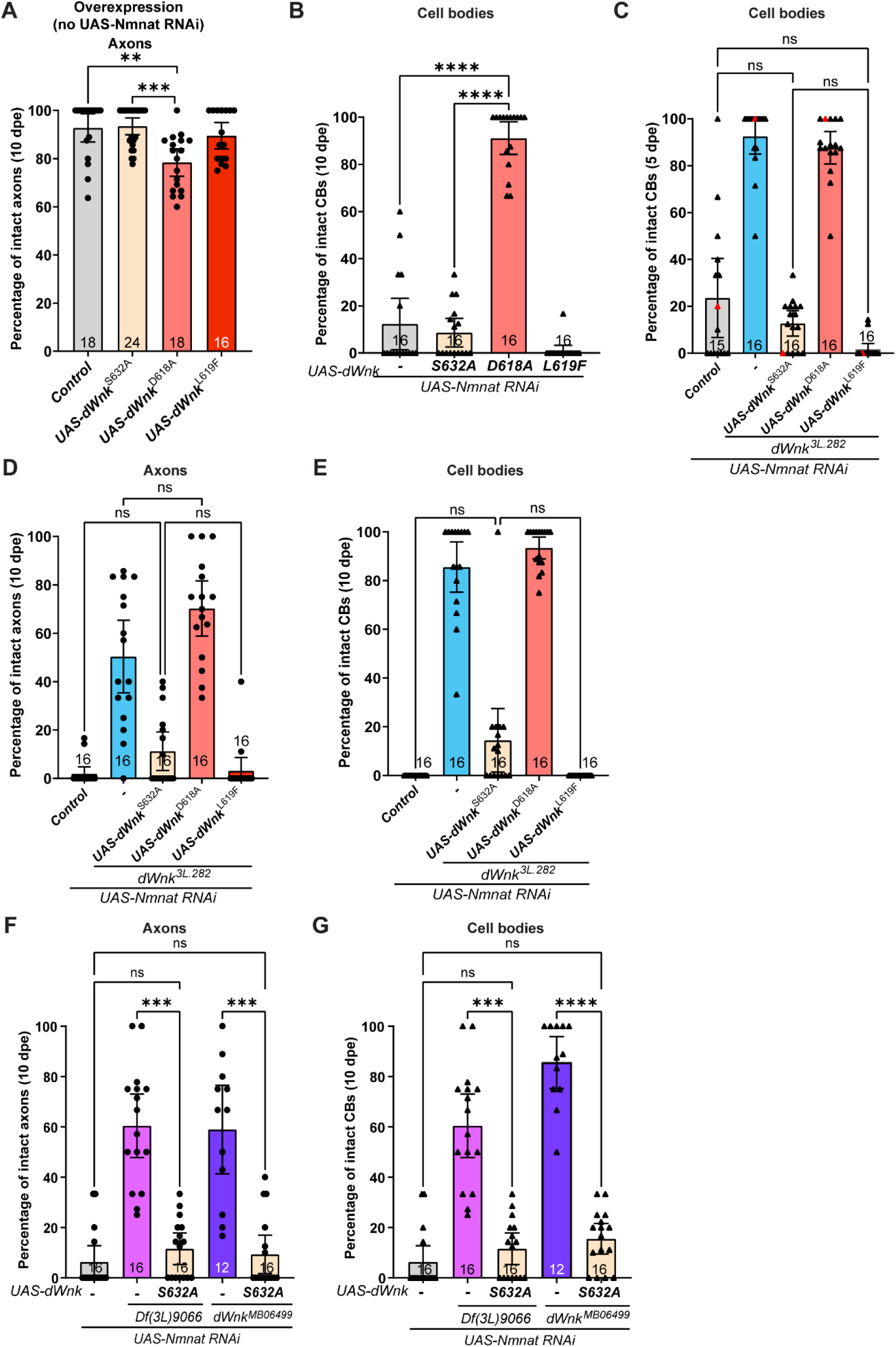
Related to Figure 2. **(A)** Overexpression of dWnk^S632A^ and dWnk^L619F^ without *dNmnat KD* does not cause neurodegeneration. Overexpression of dWnk^D618A^ without *dNmnat KD* slightly reduces neuronal viability. The results are plotted as mean ± 95% CI and analyzed using Kruskal-Wallis tests with post-hoc Dunn’s multiple comparison test. All comparisons are listed in the Supplementary table **(B)** Quantifications showing the neuronal cell body protection provided by the expression of dWnk^D618A^ against *dNmnat KD* (10 dpe, related to Fig. 2B). The results are plotted as mean ± 95% CI and analyzed using Kruskal-Wallis tests with post-hoc Dunn’s multiple comparison test. All comparisons are listed in the Supplementary table 2. **(C)** Re-supplying the kinase-dead dWnk^D618A^ cannot erase the cell body protection provided by the *dWnk^3L.282^* allele (5 dpe). Red triangles correspond to representative images in Fig. 2C. The results are plotted as mean ± 95% CI and analyzed using Kruskal-Wallis tests with post-hoc Dunn’s multiple comparison test. All comparisons are listed in the Supplementary table 2. **(D-E)** Expression of the kinase-dead dWnk^D618A^ does not erase the axonal (D) and cell body (E) protection provided by the *dWnk^3L.282^*allele, even at 10 dpe. In contrast, expression of either dWnk^S632A^ or dWnk^L619F^ completely erases the axonal (D) and cell body (E) protection provided by *dWnk^3L.282^*by 10 dpe. The results are plotted as mean ± 95% CI and analyzed using Kruskal-Wallis tests with post-hoc Dunn’s multiple comparison test. All comparisons are listed in the Supplementary table 2. **(F-G)** Expression of auto-activation impaired dWnk^S632A^ completely erased the axon (F) and cell body (G) protection provided by either *dWnk^MB06499^* or *Df(3L)Exel9066* by 10 dpe. The results are plotted as mean ± 95% CI and analyzed using Kruskal-Wallis tests with post-hoc Dunn’s multiple comparison test. All comparisons are listed in the Supplementary table 2.

**Figure S3.**
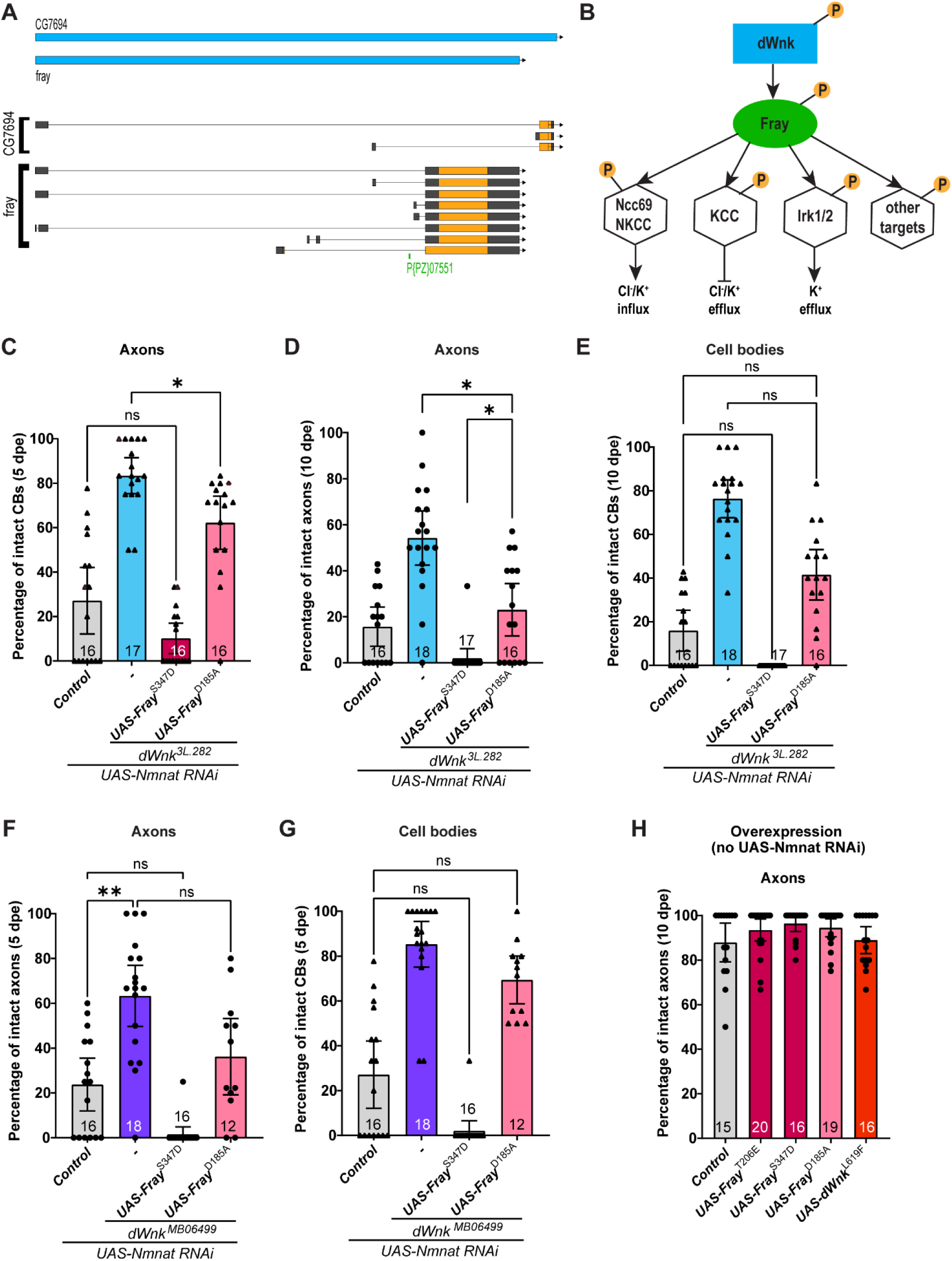
Related to Figure 3. **(A)** Schematic representation of the *fray* locus, indicating the *fray^07551^* allele used in this study. **(B)** The dWnk pathway: dWnk is activated by auto-phosphorylation. Fray is then phosphorylated by dWnk at the T-loop residue T206 and the S-motif residue S347. Activated Fray subsequently phosphorylates downstream targets. **(C)** Expression of the phospho-mimicking form of Fray, Fray^S347D^, abolishes cell body protection provided by *dWnk^3L.282^* against dNmnat depletion (5 dpe). Expression of the kinase-impaired form of Fray, Fray^D185A^, in *dWnk^3L.282^*neurons delays *dNmnat KD*-induced cell body loss (5 dpe). The results are plotted as mean ± 95% CI and analyzed using Kruskal-Wallis tests with post-hoc Dunn’s multiple comparison test. All comparisons are listed in the Supplementary table 2. **(D-E)** Expression of Fray^S347D^ completely erases the axonal (D) and cell body (E) protection provided by *dWnk^3L.282^* by 10 dpe. Expression of Fray^D185A^ in *dWnk^3L.282^* neurons significantly delays axonal (D) and cell body (E) degeneration at 10 dpe. The results are plotted as mean ± 95% CI and analyzed using Kruskal-Wallis tests with post-hoc Dunn’s multiple comparison test. All comparisons are listed in the Supplementary table 2. **(F-G)** Expression of Fray^S347D^ completely erases the axonal (F) and cell body (G) protection provided by *dWnk^MB06499^*by 5 dpe. Expression Fray^D185A^, in *dWnk^MB06499^* neurons significantly delays axonal (F) and cell body (G) degeneration at 5 dpe. The results are plotted as mean ± 95% CI and analyzed using Kruskal-Wallis tests with post-hoc Dunn’s multiple comparison test. All comparisons are listed in the Supplementary table 2. **(H)** Overexpression of Fray^T206E^, Fray^S347D^, Fray^D185A^ and dWnk^L619F^ without *dNmnat KD* does not cause neurodegeneration by 10 dpe. The results are plotted as mean ± 95% CI. Data analysis is provided in the Supplementary table 2.

**Figure S4.**
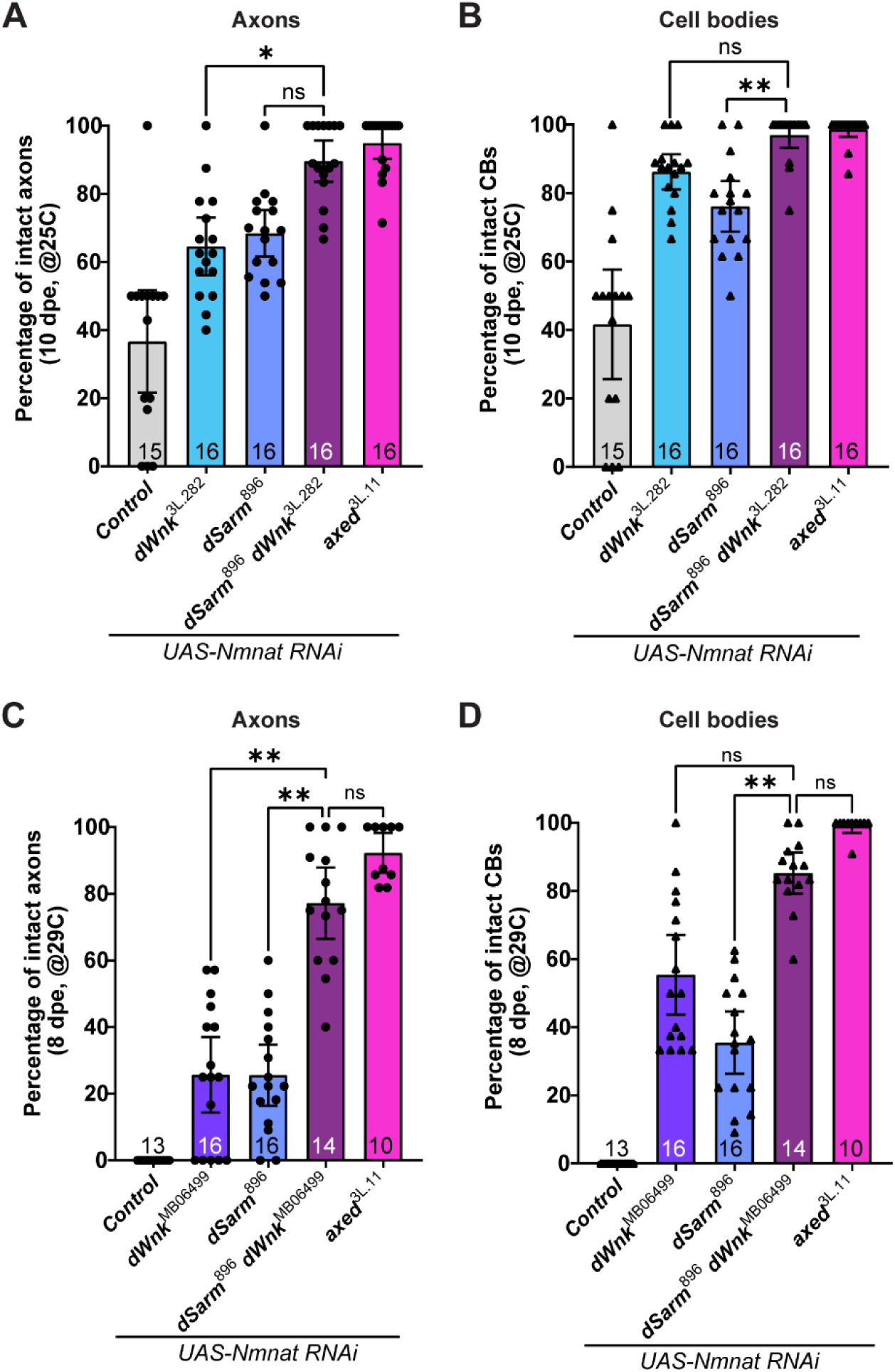
Related to Figure 4. **(A-B)** The protective effect of the *dSarm^896^,dWnk^3L.282^* double mutant is additive compared to the single mutants, and tantamount to *axed^3L.11^* protection levels in a standard *dNmnat KD* experiment (25°C). The results are plotted as mean ± 95% CI and analyzed using Kruskal-Wallis tests with post-hoc Dunn’s multiple comparison test. All comparisons are listed in the Supplementary table **(C-D)** *dWnk^MB06499^* and *dSarm^896^* partially protect axons (C) and neuronal cell bodies (D) from strong dNmnat depletion (29°C). *dSarm^896^,dWnk^MB06499^* double mutant axons (C) and cell bodies (D) are protected to the same extent as *axed KO*. The results are plotted as mean ± 95% CI and analyzed using Kruskal-Wallis tests with post-hoc Dunn’s multiple comparison test. All comparisons are listed in the Supplementary table 2.

**Figure S5.**
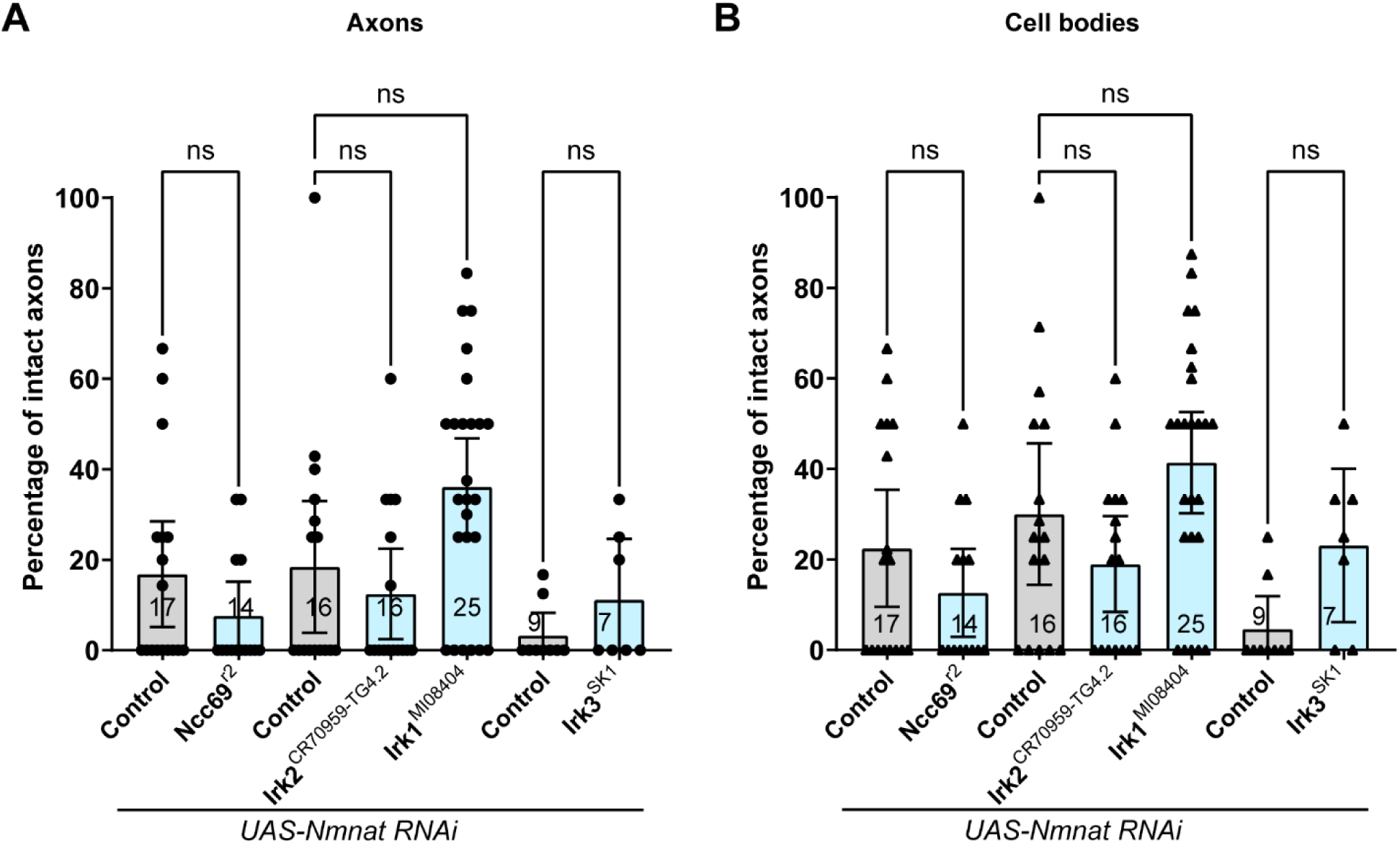
Related to Figure 5. **(A-B)** Knocking out known *Fray* downstream targets individually does not confer neuroprotection against *dNmnat* depletion. Results are presented as mean ± 95% CI and were analyzed using Kruskal-Wallis tests followed by Dunn’s multiple comparison post-hoc test. All comparisons are provided in Supplementary Table 2.

