## Supplementary Tables 1-3 for "An Ionic Sensor acts in Parallel to dSarm to Promote Neurodegeneration"

### Supplementary Table 1

| Kruskal-Wallis tests with post-hoc Dunn's multiple comparisons test | Mean rank diff. | Significant? | Summary | Adjusted P Value |
|---|---|---|---|---|
| **Figure 1C** | | | | |
| Control vs. $dWnk^{3L.282}$ | -33.63 | Yes | *** | 0.0003 |
| Control vs. Df(3L)Exel9066 | -37.02 | Yes | **** | <0.0001 |
| Control vs. $dWnk^{MB06499}$ | -36.63 | Yes | *** | 0.0003 |
| Control vs. $dWnk^{3L.282}$ + UAS-$dWnk^{WT}$ | 3.143 | No | ns | >0.9999 |
| $dWnk^{3L.282}$ vs. $dWnk^{3L.282}$ + UAS-$dWnk^{WT}$ | 36.77 | Yes | *** | 0.0002 |
| **Figure 1D** | | | | |
| Control vs. $dWnk^{3L.282}$ | -33.33 | Yes | *** | 0.0004 |
| Control vs. Df(3L)Exel9066 | -35.95 | Yes | *** | 0.0001 |
| Control vs. $dWnk^{MB06499}$ | -45.68 | Yes | **** | <0.0001 |
| Control vs. $dWnk^{3L.282}$ + UAS-$dWnk^{WT}$ | 4.305 | No | ns | >0.9999 |
| $dWnk^{3L.282}$ vs. $dWnk^{3L.282}$ + UAS-$dWnk^{WT}$ | 37.63 | Yes | *** | 0.0001 |
| **Figure 2B** | | | | |
| - vs. S632A | -4.781 | No | ns | >0.9999 |
| - vs. D618A | -28.03 | Yes | **** | <0.0001 |
| - vs. L619F | 7.063 | No | ns | >0.9999 |
| S632A vs. D618A | -23.25 | Yes | *** | 0.0005 |
| S632A vs. L619F | 11.84 | No | ns | 0.2705 |
| D618A vs. L619F | 35.09 | Yes | **** | <0.0001 |
| **Figure 2D** | | | | |
| Control vs. $dWnk^{3L.282}$ | -31.52 | Yes | ** | 0.0011 |
| Control vs. $dWnk^{3L.282}$ + UAS-$dWnk^{S632A}$ | -3.898 | No | ns | >0.9999 |
| Control vs. $dWnk^{3L.282}$ + UAS-$dWnk^{D618A}$ | -33.65 | Yes | *** | 0.0004 |
| Control vs. $dWnk^{3L.282}$ + UAS-$dWnk^{L619F}$ | 10.48 | No | ns | >0.9999 |
| $dWnk^{3L.282}$ vs. $dWnk^{3L.282}$ + UAS-$dWnk^{S632A}$ | 27.63 | Yes | ** | 0.0058 |
| $dWnk^{3L.282}$ vs. $dWnk^{3L.282}$ + UAS-$dWnk^{D618A}$ | -2.125 | No | ns | >0.9999 |
| $dWnk^{3L.282}$ vs. $dWnk^{3L.282}$ + UAS-$dWnk^{L619F}$ | 42 | Yes | **** | <0.0001 |
| $dWnk^{3L.282}$ + UAS-$dWnk^{S632A}$ vs. $dWnk^{3L.282}$ + UAS-$dWnk^{D618A}$ | -29.75 | Yes | ** | 0.0021 |
| $dWnk^{3L.282}$ + UAS-$dWnk^{S632A}$ vs. $dWnk^{3L.282}$ + UAS-$dWnk^{L619F}$ | 14.38 | No | ns | 0.7328 |
| $dWnk^{3L.282}$ + UAS-$dWnk^{D618A}$ vs. $dWnk^{3L.282}$ + UAS-$dWnk^{L619F}$ | 44.13 | Yes | **** | <0.0001 |
| **Figure 3C** | | | | |
| Control vs. $fray^{07551}$ | -24.38 | Yes | ** | 0.0012 |
| Control vs. $fray^{\Delta}$ | -23.91 | Yes | ** | 0.0016 |
| Control vs. $fray^{K67M}$ | -15.09 | No | ns | 0.1275 |
| $fray^{07551}$ vs. $fray^{\Delta}$ | 0.4688 | No | ns | >0.9999 |
| $fray^{07551}$ vs. $fray^{K67M}$ | 9.281 | No | ns | 0.9399 |
| $fray^{\Delta}$ vs. $fray^{K67M}$ | 8.813 | No | ns | >0.9999 |

| | | | | |
|---|---|---|---|---|
| **Figure 3D** | | | | |
| Control vs. fray$^{07551}$ | -42.09 | Yes | *** | 0.0005 |
| fray$^{07551}$ vs. fray$^{07551}$ + UAS-Fray$^{T206E}$ | 58.77 | Yes | **** | <0.0001 |
| fray$^{07551}$ vs. fray$^{07551}$ + UAS-Fray$^{S347D}$ | 44.27 | Yes | *** | 0.0001 |
| fray$^{07551}$ vs. fray$^{07551}$ + UAS-Fray$^{D185A}$ | 10.33 | No | ns | >0.9999 |
| fray$^{07551}$ vs. fray$^{07551}$ + UAS-dWnk$^{L619F}$ | 4.892 | No | ns | >0.9999 |
| Control vs. fray$^{07551}$ + UAS-Fray$^{T206E}$ | 16.67 | No | ns | >0.9999 |
| Control vs. fray$^{07551}$ + UAS-Fray$^{S347D}$ | 2.180 | No | ns | >0.9999 |
| Control vs. fray$^{07551}$ + UAS-Fray$^{D185A}$ | -31.76 | Yes | * | 0.0195 |
| Control vs. fray$^{07551}$ + UAS-dWnk$^{L619F}$ | -37.20 | Yes | ** | 0.0027 |
| fray$^{07551}$ + UAS-Fray$^{T206E}$ vs. fray$^{07551}$ + UAS-Fray$^{S347D}$ | -14.49 | No | ns | >0.9999 |
| fray$^{07551}$ + UAS-Fray$^{T206E}$ vs. fray$^{07551}$ + UAS-Fray$^{D185A}$ | -48.43 | Yes | **** | <0.0001 |
| fray$^{07551}$ + UAS-Fray$^{S347D}$ vs. fray$^{07551}$ + UAS-Fray$^{D185A}$ | -33.94 | Yes | ** | 0.0075 |
| fray$^{07551}$ + UAS-dWnk$^{L619F}$ vs. fray$^{07551}$ + UAS-Fray$^{D185A}$ | 5.441 | No | ns | >0.9999 |
| fray$^{07551}$ + UAS-dWnk$^{L619F}$ vs. fray$^{07551}$ + UAS-Fray$^{T206E}$ | 53.88 | Yes | **** | <0.0001 |
| fray$^{07551}$ + UAS-dWnk$^{L619F}$ vs. fray$^{07551}$ + UAS-Fray$^{S347D}$ | 39.38 | Yes | ** | 0.0011 |
| **Figure 3E** | | | | |
| Control vs. fray$^{07551}$ | -45.16 | Yes | *** | 0.0001 |
| fray$^{07551}$ vs. fray$^{07551}$ + UAS-Fray$^{T206E}$ | 66.67 | Yes | **** | <0.0001 |
| fray$^{07551}$ vs. fray$^{07551}$ + UAS-Fray$^{S347D}$ | 50.07 | Yes | **** | <0.0001 |
| fray$^{07551}$ vs. fray$^{07551}$ + UAS-Fray$^{D185A}$ | 15.72 | No | ns | >0.9999 |
| fray$^{07551}$ vs. fray$^{07551}$ + UAS-dWnk$^{L619F}$ | 3.809 | No | ns | >0.9999 |
| Control vs. fray$^{07551}$ + UAS-Fray$^{T206E}$ | 21.51 | No | ns | 0.3510 |
| Control vs. fray$^{07551}$ + UAS-Fray$^{S347D}$ | 4.917 | No | ns | >0.9999 |
| Control vs. fray$^{07551}$ + UAS-Fray$^{D185A}$ | -29.44 | Yes | * | 0.0390 |
| Control vs. fray$^{07551}$ + UAS-dWnk$^{L619F}$ | -41.35 | Yes | *** | 0.0004 |
| fray$^{07551}$ + UAS-Fray$^{T206E}$ vs. fray$^{07551}$ + UAS-Fray$^{S347D}$ | -16.59 | No | ns | >0.9999 |
| fray$^{07551}$ + UAS-Fray$^{T206E}$ vs. fray$^{07551}$ + UAS-Fray$^{D185A}$ | -50.95 | Yes | **** | <0.0001 |
| fray$^{07551}$ + UAS-Fray$^{S347D}$ vs. fray$^{07551}$ + UAS-Fray$^{D185A}$ | -34.35 | Yes | ** | 0.0058 |
| fray$^{07551}$ + UAS-dWnk$^{L619F}$ vs. fray$^{07551}$ + UAS-Fray$^{T206E}$ | 62.86 | Yes | **** | <0.0001 |
| fray$^{07551}$ + UAS-dWnk$^{L619F}$ vs. fray$^{07551}$ + UAS-Fray$^{S347D}$ | 46.26 | Yes | **** | <0.0001 |
| fray$^{07551}$ + UAS-dWnk$^{L619F}$ vs. fray$^{07551}$ + UAS-Fray$^{D185A}$ | 11.91 | No | ns | >0.9999 |
| **Figure 3F** | | | | |
| Control vs. dWnk$^{3L.282}$ | -22.66 | Yes | ** | 0.0030 |
| Control vs. dWnk$^{3L.282}$ + UAS-Fray$^{S347D}$ | 10.84 | No | ns | 0.6069 |
| Control vs. dWnk$^{3L.282}$ + UAS-Fray$^{D185A}$ | -9.750 | No | ns | 0.8430 |
| dWnk$^{3L.282}$ vs. dWnk$^{3L.282}$ + UAS-Fray$^{S347D}$ | 33.50 | Yes | **** | <0.0001 |

| | | | | |
|---|---|---|---|---|
| dWnk$^{3L.282}$ vs. dWnk$^{3L.282}$ + UAS-Fray$^{D185A}$ | 12.91 | No | ns | 0.2859 |
| dWnk$^{3L.282}$ + UAS-Fray$^{S347D}$ vs. dWnk$^{3L.282}$ + UAS-Fray$^{D185A}$ | -20.59 | Yes | * | 0.0111 |
| **Figure 4B** | | | | |
| Control vs. dWnk$^{3L.282}$ | -16.56 | No | ns | 0.4195 |
| Control vs. dSarm$^{896}$ | -25.44 | Yes | * | 0.0178 |
| Control vs. dSarm$^{896}$ dWnk$^{3L.282}$ | -47.66 | Yes | **** | <0.0001 |
| Control vs. axed$^{3L.11}$ | -53.00 | Yes | **** | <0.0001 |
| dWnk$^{3L.282}$ vs. dSarm$^{896}$ | -8.875 | No | ns | >0.9999 |
| dWnk$^{3L.282}$ vs. dSarm$^{896}$ dWnk$^{3L.282}$ | -31.09 | Yes | ** | 0.0013 |
| dWnk$^{3L.282}$ vs. axed$^{3L.11}$ | -36.44 | Yes | **** | <0.0001 |
| dSarm$^{896}$ vs. dSarm$^{896}$ dWnk$^{3L.282}$ | -22.22 | No | ns | 0.0636 |
| dSarm$^{896}$ vs. axed$^{3L.11}$ | -27.56 | Yes | ** | 0.0071 |
| dSarm$^{896}$ dWnk$^{3L.282}$ vs. axed$^{3L.11}$ | -5.344 | No | ns | >0.9999 |
| **Figure 4C** | | | | |
| Control vs. dWnk$^{3L.282}$ | -33.38 | Yes | *** | 0.0004 |
| Control vs. dSarm$^{896}$ | -19.69 | No | ns | 0.1465 |
| Control vs. dSarm$^{896}$ dWnk$^{3L.282}$ | -49.75 | Yes | **** | <0.0001 |
| Control vs. axed$^{3L.11}$ | -48.91 | Yes | **** | <0.0001 |
| dWnk$^{3L.282}$ vs. dSarm$^{896}$ | 13.69 | No | ns | 0.8968 |
| dWnk$^{3L.282}$ vs. dSarm$^{896}$ dWnk$^{3L.282}$ | -16.38 | No | ns | 0.4232 |
| dWnk$^{3L.282}$ vs. axed$^{3L.11}$ | -15.53 | No | ns | 0.5414 |
| dSarm$^{896}$ vs. dSarm$^{896}$ dWnk$^{3L.282}$ | -30.06 | Yes | ** | 0.0019 |
| dSarm$^{896}$ vs. axed$^{3L.11}$ | -29.22 | Yes | ** | 0.0029 |
| dSarm$^{896}$ dWnk$^{3L.282}$ vs. axed$^{3L.11}$ | 0.8438 | No | ns | >0.9999 |
| **Figure 4D** | | | | |
| Control vs. dWnk$^{3L.282}$ | 10.05 | No | ns | 0.1348 |
| Control vs. fray$^{07551}$ | -4.371 | No | ns | 0.7550 |
| **Figure 4E** | | | | |
| Control vs. dSarm$^{896}$ | -35.66 | Yes | **** | <0.0001 |
| Control vs. dSarm$^{896}$ + UAS-dWnk$^{L619F}$ | -14.22 | No | ns | 0.1732 |
| Control vs. dSarm$^{896}$ + UAS-dWnk$^{S632A}$ | -36.88 | Yes | **** | <0.0001 |
| dSarm$^{896}$ vs. dSarm$^{896}$ + UAS-dWnk$^{L619F}$ | 21.44 | Yes | ** | 0.0059 |
| dSarm$^{896}$ vs. dSarm$^{896}$ + UAS-dWnk$^{S632A}$ | -1.219 | No | ns | >0.9999 |
| dSarm$^{896}$ + UAS-dWnk$^{L619F}$ vs. dSarm$^{896}$ + UAS-dWnk$^{S632A}$ | -22.66 | Yes | ** | 0.0030 |
| **Figure 4F** | | | | |
| Control vs. dSarm$^{896}$ | -27.53 | Yes | *** | 0.0001 |
| Control vs. dSarm$^{896}$ + UAS-dWnk$^{L619F}$ | -31.91 | Yes | **** | <0.0001 |
| Control vs. dSarm$^{896}$ + UAS-dWnk$^{S632A}$ | -36.56 | Yes | **** | <0.0001 |
| dSarm$^{896}$ vs. dSarm$^{896}$ + UAS-dWnk$^{L619F}$ | -4.375 | No | ns | >0.9999 |
| dSarm$^{896}$ vs. dSarm$^{896}$ + UAS-dWnk$^{S632A}$ | -9.031 | No | ns | >0.9999 |
| dSarm$^{896}$ + UAS-dWnk$^{L619F}$ vs. dSarm$^{896}$ + UAS-dWnk$^{S632A}$ | -4.656 | No | ns | >0.9999 |

| | | | | |
|---|---|---|---|---|
| **Figure 4G** | | | | |
| Control vs. $dSarm^{896}$ | -33.94 | Yes | **** | <0.0001 |
| Control vs. $dSarm^{896}$ + UAS-$Fray^{S347D}$ | -8.927 | No | ns | 0.9683 |
| Control vs. $dSarm^{896}$ + UAS-$Fray^{D185A}$ | -26.97 | Yes | *** | 0.0001 |
| $dSarm^{896}$ vs. $dSarm^{896}$ + UAS-$Fray^{S347D}$ | 25.01 | Yes | *** | 0.0005 |
| $dSarm^{896}$ vs. $dSarm^{896}$ + UAS-$Fray^{D185A}$ | 6.969 | No | ns | >0.9999 |
| $dSarm^{896}$ + UAS-$Fray^{S347D}$ vs. $dSarm^{896}$ + UAS-$Fray^{D185A}$ | -18.04 | Yes | * | 0.0279 |
| **Figure 4H** | | | | |
| Control vs. $dSarm^{896}$ | -34.31 | Yes | **** | <0.0001 |
| Control vs. $dSarm^{896}$ + UAS-$Fray^{S347D}$ | -14.45 | No | ns | 0.1535 |
| Control vs. $dSarm^{896}$ + UAS-$Fray^{D185A}$ | -25.72 | Yes | *** | 0.0003 |
| $dSarm^{896}$ vs. $dSarm^{896}$ + UAS-$Fray^{S347D}$ | 19.86 | Yes | * | 0.0130 |
| $dSarm^{896}$ vs. $dSarm^{896}$ + UAS-$Fray^{D185A}$ | 8.594 | No | ns | >0.9999 |
| $dSarm^{896}$ + UAS-$Fray^{S347D}$ vs. $dSarm^{896}$ + UAS-$Fray^{D185A}$ | -11.26 | No | ns | 0.4913 |
| **Figure 4I** | | | | |
| UAS-lacZ vs. UAS-$dWnk^{L619F}$ | -2.989 | No | ns | >0.9999 |
| UAS-lacZ vs. UAS-$dWnk^{S632A}$ | -7.450 | No | ns | >0.9999 |
| UAS-lacZ vs. UAS-$Fray^{S347D}$ | -4.997 | No | ns | >0.9999 |
| UAS-lacZ vs. UAS-$Fray^{D185A}$ | 1.503 | No | ns | >0.9999 |
| UAS-$dWnk^{L619F}$ vs. UAS-$dWnk^{S632A}$ | -4.461 | No | ns | >0.9999 |
| UAS-$dWnk^{L619F}$ vs. UAS-$Fray^{S347D}$ | -2.008 | No | ns | >0.9999 |
| UAS-$dWnk^{L619F}$ vs. UAS-$Fray^{D185A}$ | 4.492 | No | ns | >0.9999 |
| UAS-$dWnk^{S632A}$ vs. UAS-$Fray^{S347D}$ | 2.453 | No | ns | >0.9999 |
| UAS-$dWnk^{S632A}$ vs. UAS-$Fray^{D185A}$ | 8.953 | No | ns | >0.9999 |
| UAS-$Fray^{S347D}$ vs. UAS-$Fray^{D185A}$ | 6.500 | No | ns | >0.9999 |
| **Figure 5A** | | | | |
| Control vs. $axed^{3L.11}$ | -24.22 | Yes | **** | <0.0001 |
| Control vs. $axed^{3L.11}$ + UAS-$Fray^{S347D}$ | -23.78 | Yes | **** | <0.0001 |
| $axed^{3L.11}$ vs. $axed^{3L.11}$ + UAS-$Fray^{S347D}$ | 0.4375 | No | ns | >0.9999 |

| **Two-way ANOVA, Šídák's multiple comparisons test** | **Predicted (LS) mean diff.** | **95.00% CI of diff.** | **Below threshold?** | **Summary** | **Adjusted P Value** |
|---|---|---|---|---|---|
| **Axons – Cell Bodies** | | | | | |
| Control | -0.05556 | -1.220 to 1.109 | No | ns | >0.9999 |
| $dWnk^{3L.282}$ | 0.09091 | -0.9624 to 1.144 | No | ns | >0.9999 |
| $dSarm^{E1170A}$ | 0.375 | -1.372 to 2.122 | No | ns | 0.9934 |
| $dSarm^{E1170A}dWnk^{3L.282}$ | 0.3846 | -0.9857 to 1.755 | No | ns | 0.9737 |
| $dSarm^{896}$ | 3.167 | 1.150 to 5.184 | Yes | *** | 0.0003 |
| $dSarm^{896}$ + UAS-$Fray^{S347D}$ | 2.167 | 0.1497 to 4.184 | Yes | * | 0.0284 |

### Supplementary Table 2

| Figure S1I | |
|---|---|
| **Mann Whitney test** | |
| P value | 0.2004 |
| Exact or approximate P value? | Exact |
| P value summary | ns |
| Significantly different (P < 0.05)? | No |
| One- or two-tailed P value? | Two-tailed |
| Sum of ranks in column A,B | 304.5, 256.5 |
| Mann-Whitney U | 103.5 |

| **Kruskal-Wallis tests with post-hoc Dunn's multiple comparisons test** | **Mean rank diff.** | **Significant?** | **Summary** | **Adjusted P Value** |
|---|---|---|---|---|
| **Figure S1D** | | | | |
| 1 dpe vs. 5 dpe | 3.531 | No | ns | >0.9999 |
| 1 dpe vs. 10 dpe | 16.99 | Yes | ** | 0.0011 |
| 5 dpe vs. 10 dpe | 13.46 | Yes | * | 0.0140 |
| **Figure S1E** | | | | |
| 1 dpe vs. 5 dpe | 11.31 | No | ns | 0.0715 |
| 1 dpe vs. 10 dpe | 24.26 | Yes | **** | <0.0001 |
| 5 dpe vs. 10 dpe | 12.95 | Yes | * | 0.0307 |
| **Figure S1F** | | | | |
| Control (-) vs. Control (+) | 11.23 | No | ns | >0.9999 |
| $dWnk^{3L.1541}$ (-) vs. $dWnk^{3L.1541}$ (+) | 64.66 | Yes | **** | <0.0001 |
| $dWnk^{MB06499}$ (-) vs. $dWnk^{MB06499}$ (+) | 66.44 | Yes | **** | <0.0001 |
| Df(3L)9066 (-) vs. Df(3L)9066 (+) | 61.75 | Yes | **** | <0.0001 |
| **Figure S1G** | | | | |
| Control vs. $dWnk^{F1183}$ | -6.548 | No | ns | 0.3178 |
| Control vs. $dWnk^{G1286}$ | -7.292 | No | ns | 0.3315 |
| $dWnk^{F1183}$ vs. $dWnk^{G1286}$ | -0.744 | No | ns | >0.9999 |
| **Figure S1H** | | | | |
| Control vs. $dWnk^{F1183}$ | -9.929 | No | ns | >0.9999 |
| Control vs. $dWnk^{G1286}$ | -8.414 | No | ns | >0.9999 |
| Control vs. $dWnk^{3L.282}$ | -21.85 | No | ns | 0.0995 |
| Control vs. $dWnk^{MB06499}$ | -3.019 | No | ns | >0.9999 |
| Control vs. Df(3L)Exel9066 | 14.69 | No | ns | 0.7729 |
| $dWnk^{F1183}$ vs. $dWnk^{G1286}$ | 1.514 | No | ns | >0.9999 |
| $dWnk^{F1183}$ vs. $dWnk^{3L.282}$ | -11.92 | No | ns | >0.9999 |
| $dWnk^{F1183}$ vs. $dWnk^{MB06499}$ | 6.91 | No | ns | >0.9999 |
| $dWnk^{F1183}$ vs. Df(3L)Exel9066 | 24.62 | No | ns | 0.0703 |
| $dWnk^{G1286}$ vs. $dWnk^{3L.282}$ | -13.44 | No | ns | >0.9999 |
| $dWnk^{G1286}$ vs. $dWnk^{MB06499}$ | 5.396 | No | ns | >0.9999 |
| $dWnk^{G1286}$ vs. Df(3L)Exel9066 | 23.1 | No | ns | 0.2989 |
| $dWnk^{3L.282}$ vs. $dWnk^{MB06499}$ | 18.83 | No | ns | 0.6465 |
| $dWnk^{3L.282}$ vs. Df(3L)Exel9066 | 36.54 | Yes | ** | 0.0013 |

| | | | | |
|---|---|---|---|---|
| dWnk$^{MB06499}$ vs. Df(3L)Exel9066 | 17.71 | No | ns | 0.6911 |
| **Figure S2A** | | | | |
| Control vs. UAS-dWnk$^{S632A}$ | -0.02083 | No | ns | >0.9999 |
| Control vs. UAS-dWnk$^{D618A}$ | 24.92 | Yes | ** | 0.0023 |
| Control vs. UAS-dWnk$^{L619F}$ | 8.021 | No | ns | >0.9999 |
| UAS-dWnk$^{S632A}$ vs. UAS-dWnk$^{D618A}$ | 24.94 | Yes | *** | 0.0009 |
| UAS-dWnk$^{S632A}$ vs. UAS-dWnk$^{L619F}$ | 8.042 | No | ns | >0.9999 |
| UAS-dWnk$^{D618A}$ vs. UAS-dWnk$^{L619F}$ | -16.9 | No | ns | 0.117 |
| **Figure S2B** | | | | |
| - vs. S632A | -1.156 | No | ns | >0.9999 |
| - vs. D618A | -30 | Yes | **** | <0.0001 |
| - vs. L619F | 7.156 | No | ns | >0.9999 |
| S632A vs. D618A | -28.84 | Yes | **** | <0.0001 |
| S632A vs. L619F | 8.313 | No | ns | 0.9983 |
| D618A vs. L619F | 37.16 | Yes | **** | <0.0001 |
| **Figure S2C** | | | | |
| Control vs. dWnk$^{3L.282}$ | -35.03 | Yes | *** | 0.0001 |
| Control vs. dWnk$^{3L.282}$ + UAS-dWnk$^{S632A}$ | 1.85 | No | ns | >0.9999 |
| Control vs. dWnk$^{3L.282}$ + UAS-dWnk$^{D618A}$ | -30.09 | Yes | ** | 0.0019 |
| Control vs. dWnk$^{3L.282}$ + UAS-dWnk$^{L619F}$ | 14.38 | No | ns | 0.7421 |
| dWnk$^{3L.282}$ vs. dWnk$^{3L.282}$ + UAS-dWnk$^{S632A}$ | 36.88 | Yes | **** | <0.0001 |
| dWnk$^{3L.282}$ vs. dWnk$^{3L.282}$ + UAS-dWnk$^{D618A}$ | 4.938 | No | ns | >0.9999 |
| dWnk$^{3L.282}$ vs. dWnk$^{3L.282}$ + UAS-dWnk$^{L619F}$ | 49.41 | Yes | **** | <0.0001 |
| dWnk$^{3L.282}$ + UAS-dWnk$^{S632A}$ vs. dWnk$^{3L.282}$ + UAS-dWnk$^{D618A}$ | -31.94 | Yes | *** | 0.0006 |
| dWnk$^{3L.282}$ + UAS-dWnk$^{S632A}$ vs. dWnk$^{3L.282}$ + UAS-dWnk$^{L619F}$ | 12.53 | No | ns | >0.9999 |
| dWnk$^{3L.282}$ + UAS-dWnk$^{D618A}$ vs. dWnk$^{3L.282}$ + UAS-dWnk$^{L619F}$ | 44.47 | Yes | **** | <0.0001 |
| **Figure S2D** | | | | |
| Control vs. dWnk$^{3L.282}$ | -35.41 | Yes | **** | <0.0001 |
| Control vs. dWnk$^{3L.282}$ + UAS-dWnk$^{S632A}$ | -9.563 | No | ns | >0.9999 |
| Control vs. dWnk$^{3L.282}$ + UAS-dWnk$^{D618A}$ | -44.94 | Yes | **** | <0.0001 |
| Control vs. dWnk$^{3L.282}$ + UAS-dWnk$^{L619F}$ | -0.7188 | No | ns | >0.9999 |
| dWnk$^{3L.282}$ vs. dWnk$^{3L.282}$ + UAS-dWnk$^{S632A}$ | 25.84 | Yes | ** | 0.0087 |
| dWnk$^{3L.282}$ vs. dWnk$^{3L.282}$ + UAS-dWnk$^{D618A}$ | -9.531 | No | ns | >0.9999 |
| dWnk$^{3L.282}$ vs. dWnk$^{3L.282}$ + UAS-dWnk$^{L619F}$ | 34.69 | Yes | **** | <0.0001 |
| dWnk$^{3L.282}$ + UAS-dWnk$^{S632A}$ vs. dWnk$^{3L.282}$ + UAS-dWnk$^{D618A}$ | -35.38 | Yes | **** | <0.0001 |
| dWnk$^{3L.282}$ + UAS-dWnk$^{S632A}$ vs. dWnk$^{3L.282}$ + UAS-dWnk$^{L619F}$ | 8.844 | No | ns | >0.9999 |
| dWnk$^{3L.282}$ + UAS-dWnk$^{D618A}$ vs. dWnk$^{3L.282}$ + UAS-dWnk$^{L619F}$ | 44.22 | Yes | **** | <0.0001 |
| **Figure S2E** | | | | |
| Control vs. dWnk$^{3L.282}$ | -42.13 | Yes | **** | <0.0001 |
| Control vs. dWnk$^{3L.282}$ + UAS-dWnk$^{S632A}$ | -14.97 | No | ns | 0.5114 |

| | | | | |
|---|---|---|---|---|
| Control vs. $dWnk^{3L.282}$ + UAS-$dWnk^{D618A}$ | -45.41 | Yes | **** | <0.0001 |
| Control vs. $dWnk^{3L.282}$ + UAS-$dWnk^{L619F}$ | 0 | No | ns | >0.9999 |
| $dWnk^{3L.282}$ vs. $dWnk^{3L.282}$ + UAS-$dWnk^{S632A}$ | 27.16 | Yes | ** | 0.004 |
| $dWnk^{3L.282}$ vs. $dWnk^{3L.282}$ + UAS-$dWnk^{D618A}$ | -3.281 | No | ns | >0.9999 |
| $dWnk^{3L.282}$ vs. $dWnk^{3L.282}$ + UAS-$dWnk^{L619F}$ | 42.13 | Yes | **** | <0.0001 |
| $dWnk^{3L.282}$ + UAS-$dWnk^{S632A}$ vs. $dWnk^{3L.282}$ + UAS-$dWnk^{D618A}$ | -30.44 | Yes | *** | 0.0007 |
| $dWnk^{3L.282}$ + UAS-$dWnk^{S632A}$ vs. $dWnk^{3L.282}$ + UAS-$dWnk^{L619F}$ | 14.97 | No | ns | 0.5114 |
| $dWnk^{3L.282}$ + UAS-$dWnk^{D618A}$ vs. $dWnk^{3L.282}$ + UAS-$dWnk^{L619F}$ | 45.41 | Yes | **** | <0.0001 |
| **Figure S2F** | | | | |
| Ctrl vs. Df(3L)9066 (-) | -39.09 | Yes | **** | <0.0001 |
| Ctrl vs. $dWnk^{MB06499}$ (-) | -37.5 | Yes | **** | <0.0001 |
| Ctrl vs. Df(3L)9066 (+) | -6.688 | No | ns | >0.9999 |
| Ctrl vs. $dWnk^{MB06499}$ (+) | -3.281 | No | ns | >0.9999 |
| Df(3L)9066 (-) vs. Df(3L)9066 (+) | 32.41 | Yes | *** | 0.0001 |
| $dWnk^{MB06499}$ (-) vs. $dWnk^{MB06499}$ (+) | 34.22 | Yes | *** | 0.0002 |
| **Figure S2G** | | | | |
| Ctrl vs. Df(3L)9066 (-) | -38.56 | Yes | **** | <0.0001 |
| Ctrl vs. $dWnk^{MB06499}$ (-) | -48.18 | Yes | **** | <0.0001 |
| Ctrl vs. $dWnk^{MB06499}$ (+) | -10.91 | No | ns | 0.9362 |
| Ctrl vs. Df(3L)9066 (+) | -6.281 | No | ns | >0.9999 |
| Df(3L)9066 (-) vs. Df(3L)9066 (+) | 32.28 | Yes | *** | 0.0002 |
| $dWnk^{MB06499}$ (-) vs. $dWnk^{MB06499}$ (+) | 37.27 | Yes | **** | <0.0001 |
| **Figure S3C** | | | | |
| Control vs. $dWnk^{3L.282}$ | -31.30 | Yes | **** | <0.0001 |
| Control vs. $dWnk^{3L.282}$ + UAS-$Fray^{S347D}$ | 7.688 | No | ns | >0.9999 |
| Control vs. $dWnk^{3L.282}$ + UAS-$Fray^{D185A}$ | -17.59 | Yes | * | 0.0471 |
| $dWnk^{3L.282}$ vs. $dWnk^{3L.282}$ + UAS-$Fray^{S347D}$ | 38.99 | Yes | **** | <0.0001 |
| $dWnk^{3L.282}$ vs. $dWnk^{3L.282}$ + UAS-$Fray^{D185A}$ | 13.71 | No | ns | 0.2132 |
| $dWnk^{3L.282}$ + UAS-$Fray^{S347D}$ vs. $dWnk^{3L.282}$ + UAS-$Fray^{D185A}$ | -25.28 | Yes | *** | 0.0008 |
| **Figure S3D** | | | | |
| Control vs. $dWnk^{3L.282}$ | -24.66 | Yes | *** | 0.0007 |
| Control vs. $dWnk^{3L.282}$ + UAS-$Fray^{S347D}$ | 12.25 | No | ns | 0.3491 |
| Control vs. $dWnk^{3L.282}$ + UAS-$Fray^{D185A}$ | -4.906 | No | ns | >0.9999 |
| $dWnk^{3L.282}$ vs. $dWnk^{3L.282}$ + UAS-$Fray^{S347D}$ | 36.91 | Yes | **** | <0.0001 |
| $dWnk^{3L.282}$ vs. $dWnk^{3L.282}$ + UAS-$Fray^{D185A}$ | 19.75 | Yes | * | 0.0118 |
| $dWnk^{3L.282}$ + UAS-$Fray^{S347D}$ vs. $dWnk^{3L.282}$ + UAS-$Fray^{D185A}$ | -17.16 | Yes | * | 0.0479 |
| **Figure S3E** | | | | |
| Control vs. $dWnk^{3L.282}$ | -32.11 | Yes | **** | <0.0001 |
| Control vs. $dWnk^{3L.282}$ + UAS-$Fray^{S347D}$ | 10.97 | No | ns | 0.5728 |
| Control vs. $dWnk^{3L.282}$ + UAS-$Fray^{D185A}$ | -15.44 | No | ns | 0.1248 |
| $dWnk^{3L.282}$ vs. $dWnk^{3L.282}$ + UAS-$Fray^{S347D}$ | 43.08 | Yes | **** | <0.0001 |

| | | | | |
|---|---|---|---|---|
| $dWnk^{3L.282}$ vs. $dWnk^{3L.282}$ + UAS-$Fray^{D185A}$ | 16.68 | No | ns | 0.0611 |
| $dWnk^{3L.282}$ + UAS-$Fray^{S347D}$ vs. $dWnk^{3L.282}$ + UAS-$Fray^{D185A}$ | -26.41 | Yes | *** | 0.0004 |
| **Figure S3F** | | | | |
| Control vs. $dWnk^{MB06499}$ | -20.11 | Yes | ** | 0.0050 |
| Control vs. $dWnk^{MB06499}$ + UAS-$Fray^{S347D}$ | 14.38 | No | ns | 0.1210 |
| Control vs. $dWnk^{MB06499}$ + UAS-$Fray^{D185A}$ | -6.927 | No | ns | >0.9999 |
| $dWnk^{MB06499}$ vs. $dWnk^{MB06499}$ + UAS-$Fray^{S347D}$ | 34.48 | Yes | **** | <0.0001 |
| $dWnk^{MB06499}$ vs. $dWnk^{MB06499}$ + UAS-$Fray^{D185A}$ | 13.18 | No | ns | 0.2598 |
| $dWnk^{MB06499}$ + UAS-$Fray^{S347D}$ vs. $dWnk^{MB06499}$ + UAS-$Fray^{D185A}$ | -21.30 | Yes | ** | 0.0086 |
| **Figure S3G** | | | | |
| Control vs. $dWnk^{MB06499}$ | -27.00 | Yes | **** | <0.0001 |
| Control vs. $dWnk^{MB06499}$ + UAS-$Fray^{S347D}$ | 10.59 | No | ns | 0.5327 |
| Control vs. $dWnk^{MB06499}$ + UAS-$Fray^{D185A}$ | -17.54 | No | ns | 0.0545 |
| $dWnk^{MB06499}$ vs. $dWnk^{MB06499}$ + UAS-$Fray^{S347D}$ | 37.59 | Yes | **** | <0.0001 |
| $dWnk^{MB06499}$ vs. $dWnk^{MB06499}$ + UAS-$Fray^{D185A}$ | 9.458 | No | ns | 0.8967 |
| $dWnk^{MB06499}$ + UAS-$Fray^{S347D}$ vs. $dWnk^{MB06499}$ + UAS-$Fray^{D185A}$ | -28.14 | Yes | *** | 0.0002 |
| **Figure S3H** | | | | |
| Control vs. UAS-$Fray^{T206E}$ | -8.608 | No | ns | >0.9999 |
| Control vs. UAS-$Fray^{S347D}$ | -13.80 | No | ns | 0.8126 |
| Control vs. UAS-$Fray^{D185A}$ | -9.439 | No | ns | >0.9999 |
| Control vs. UAS-$dWnk^{L619F}$ | 1.729 | No | ns | >0.9999 |
| UAS-$Fray^{T206E}$ vs. UAS-$Fray^{S347D}$ | -5.194 | No | ns | >0.9999 |
| UAS-$Fray^{T206E}$ vs. UAS-$Fray^{D185A}$ | -0.8303 | No | ns | >0.9999 |
| UAS-$Fray^{T206E}$ vs. UAS-$dWnk^{L619F}$ | 10.34 | No | ns | >0.9999 |
| UAS-$Fray^{S347D}$ vs. UAS-$Fray^{D185A}$ | 4.363 | No | ns | >0.9999 |
| UAS-$Fray^{S347D}$ vs. UAS-$dWnk^{L619F}$ | 15.53 | No | ns | 0.4613 |
| UAS-$Fray^{D185A}$ vs. UAS-$dWnk^{L619F}$ | 11.17 | No | ns | >0.9999 |
| **Figure S4A** | | | | |
| Control vs. $dWnk^{3L.282}$ | -17.23 | No | ns | 0.3470 |
| Control vs. $dSarm^{896}$ | -21.29 | No | ns | 0.0906 |
| Control vs. $dSarm^{896}$ $dWnk^{3L.282}$ | -43.64 | Yes | **** | <0.0001 |
| Control vs. $axed^{3L.11}$ | -49.67 | Yes | **** | <0.0001 |
| $dWnk^{3L.282}$ vs. $dSarm^{896}$ | -4.063 | No | ns | >0.9999 |
| $dWnk^{3L.282}$ vs. $dSarm^{896}$ $dWnk^{3L.282}$ | -26.41 | Yes | * | 0.0100 |
| $dWnk^{3L.282}$ vs. $axed^{3L.11}$ | -32.44 | Yes | *** | 0.0005 |
| $dSarm^{896}$ vs. $dSarm^{896}$ $dWnk^{3L.282}$ | -22.34 | No | ns | 0.0537 |
| $dSarm^{896}$ vs. $axed^{3L.11}$ | -28.38 | Yes | ** | 0.0041 |
| $dSarm^{896}$ $dWnk^{3L.282}$ vs. $axed^{3L.11}$ | -6.031 | No | ns | >0.9999 |
| **Figure S4B** | | | | |
| Control vs. $dWnk^{3L.282}$ | -26.91 | Yes | ** | 0.0069 |
| Control vs. $dSarm^{896}$ | -16.44 | No | ns | 0.3823 |

| | | | | |
|---|---|---|---|---|
| Control vs. dSarm$^{896}$ dWnk$^{3L.282}$ | -45.23 | Yes | **** | <0.0001 |
| Control vs. axed$^{3L.11}$ | -47.69 | Yes | **** | <0.0001 |
| dWnk$^{3L.282}$ vs. dSarm$^{896}$ | 10.47 | No | ns | >0.9999 |
| dWnk$^{3L.282}$ vs. dSarm$^{896}$ dWnk$^{3L.282}$ | -18.31 | No | ns | 0.1898 |
| dWnk$^{3L.282}$ vs. axed$^{3L.11}$ | -20.78 | No | ns | 0.0776 |
| dSarm$^{896}$ vs. dSarm$^{896}$ dWnk$^{3L.282}$ | -28.78 | Yes | ** | 0.0023 |
| dSarm$^{896}$ vs. axed$^{3L.11}$ | -31.25 | Yes | *** | 0.0006 |
| dSarm$^{896}$ dWnk$^{3L.282}$ vs. axed$^{3L.11}$ | -2.469 | No | ns | >0.9999 |
| **Figure S4C** | | | | |
| Control vs. dWnk$^{MB06499}$ | -17.59 | No | ns | 0.1730 |
| Control vs. dSarm$^{896}$ | -18.34 | No | ns | 0.1307 |
| Control vs. dSarm$^{896}$ dWnk$^{MB06499}$ | -43.57 | Yes | **** | <0.0001 |
| Control vs. axed$^{3L.11}$ | -50.55 | Yes | **** | <0.0001 |
| dWnk$^{MB06499}$ vs. dSarm$^{896}$ | -0.7500 | No | ns | >0.9999 |
| dWnk$^{MB06499}$ vs. dSarm$^{896}$ dWnk$^{MB06499}$ | -25.98 | Yes | ** | 0.0034 |
| dWnk$^{MB06499}$ vs. axed$^{3L.11}$ | -32.96 | Yes | *** | 0.0004 |
| dSarm$^{896}$ vs. dSarm$^{896}$ dWnk$^{MB06499}$ | -25.23 | Yes | ** | 0.0050 |
| dSarm$^{896}$ vs. axed$^{3L.11}$ | -32.21 | Yes | *** | 0.0005 |
| dSarm$^{896}$ dWnk$^{MB06499}$ vs. axed$^{3L.11}$ | -6.979 | No | ns | >0.9999 |
| **Figure S4D** | | | | |
| Control vs. dWnk$^{MB06499}$ | -28.25 | Yes | ** | 0.0015 |
| Control vs. dSarm$^{896}$ | -18.84 | No | ns | 0.1136 |
| Control vs. dSarm$^{896}$ dWnk$^{MB06499}$ | -44.43 | Yes | **** | <0.0001 |
| Control vs. axed$^{3L.11}$ | -55.65 | Yes | **** | <0.0001 |
| dWnk$^{MB06499}$ vs. dSarm$^{896}$ | 9.406 | No | ns | >0.9999 |
| dWnk$^{MB06499}$ vs. dSarm$^{896}$ dWnk$^{MB06499}$ | -16.18 | No | ns | 0.2658 |
| dWnk$^{MB06499}$ vs. axed$^{3L.11}$ | -27.40 | Yes | ** | 0.0065 |
| dSarm$^{896}$ vs. dSarm$^{896}$ dWnk$^{MB06499}$ | -25.58 | Yes | ** | 0.0045 |
| dSarm$^{896}$ vs. axed$^{3L.11}$ | -36.81 | Yes | **** | <0.0001 |
| dSarm$^{896}$ dWnk$^{MB06499}$ vs. axed$^{3L.11}$ | -11.22 | No | ns | >0.9999 |
| **Figure S5A** | | | | |
| Control vs. Ncc69$^{r2}$ | 10.60 | No | ns | >0.9999 |
| Control vs. Irk2$^{CR70959-TG4.2}$ | 4.875 | No | ns | >0.9999 |
| Control vs. Irk1$^{MI08404}$ | -21.81 | No | ns | 0.0715 |
| Irk2$^{CR70959-TG4.2}$ vs. Irk1$^{MI08404}$ | -26.69 | Yes | * | 0.0136 |
| Control vs. Irk3$^{SK1}$ | -11.52 | No | ns | >0.9999 |
| **Figure S5B** | | | | |
| Control vs. Ncc69$^{r2}$ | 10.88 | No | ns | >0.9999 |
| Control vs. Irk2$^{CR70959-TG4.2}$ | 10.03 | No | ns | >0.9999 |
| Control vs. Irk1$^{MI08404}$ | -13.61 | No | ns | 0.7190 |
| Irk2$^{CR70959-TG4.2}$ vs. Irk1$^{MI08404}$ | -23.64 | No | ns | 0.0555 |
| Control vs. Irk3$^{SK1}$ | -25.43 | No | ns | 0.4130 |

### Supplementary Table 3: Genotypes

Figure 1:

| | |
|---|---|
| B-D 1 | w; OK371-gal4, 10xUAS-ivs-mCD8-GFP, aseFLP$^{2e}$/5xUAS-Nmnat RNAi (v32255), 5xUAS-dicer2; FRT2A FRT82B/tub-gal80 FRT2A |
| B-D 2 | w; OK371-gal4, 10xUAS-ivs-mCD8-GFP, aseFLP$^{2e}$/5xUAS-Nmnat RNAi (v32255), 5xUAS-dicer2; dWnk$^{3L.282}$ FRT2A FRT82B/tub-gal80 FRT2A |
| B-D 3 | w; OK371-gal4, 10xUAS-ivs-mCD8-GFP, aseFLP$^{2e}$/5xUAS-Nmnat RNAi (v32255), 5xUAS-dicer2; Df(3L)Exel9066 FRT2A/tub-gal80 FRT2A |
| B-D 4 | w; OK371-gal4, 10xUAS-ivs-myrTdTom, aseFLP$^{2c}$/5xUAS-Nmnat RNAi (v32255), 5xUAS-dicer2; dWnk$^{MB06499}$ FRT2A/tub-gal80 FRT2A |
| B-D 5 | w, 5xUAS-dWnk$^{WT}$/w or y; OK371-gal4, 10xUAS-ivs-mCD8-GFP, aseFLP$^{2e}$/5xUAS-Nmnat RNAi (v32255), 5xUAS-dicer2; dWnk$^{3L.282}$ FRT2A FRT82B/tub-gal80 FRT2A |

Figure 2:

| | |
|---|---|
| B1 | w; OK371-gal4, 10xUAS-ivs-mCD8-GFP, aseFLP$^{2e}$/5xUAS-Nmnat RNAi (v32255), 5xUAS-dicer2; FRT2A FRT82B/tub-gal80 FRT2A |
| B2 | w; OK371-gal4, 10xUAS-ivs-mCD8-GFP, aseFLP$^{2e}$/5xUAS-Nmnat RNAi (v32255), 5xUAS-dicer2; FRT2A 5xUAS-dWnk$^{S632A}$/tub-gal80 FRT2A |
| B3 | w; OK371-gal4, 10xUAS-ivs-mCD8-GFP, aseFLP$^{2e}$/5xUAS-Nmnat RNAi (v32255), 5xUAS-dicer2; FRT2A 5xUAS-dWnk$^{D618A}$/tub-gal80 FRT2A |
| B4 | w; OK371-gal4, 10xUAS-ivs-mCD8-GFP, aseFLP$^{2e}$/5xUAS-Nmnat RNAi (v32255), 5xUAS-dicer2; FRT2A 5xUAS-dWnk$^{L619F}$/tub-gal80 FRT2A |
| C,D 1 | w; OK371-gal4, 10xUAS-ivs-mCD8-GFP, aseFLP$^{2e}$/5xUAS-Nmnat RNAi (v32255), 5xUAS-dicer2; FRT2A FRT82B/tub-gal80 FRT2A |
| C, D 2 | w; OK371-gal4, 10xUAS-ivs-mCD8-GFP, aseFLP$^{2e}$/5xUAS-Nmnat RNAi (v32255), 5xUAS-dicer2; dWnk$^{3L.282}$ FRT2A FRT82B/tub-gal80 FRT2A |
| C,D 3 | w; OK371-gal4, 10xUAS-ivs-mCD8-GFP, aseFLP$^{2e}$/5xUAS-Nmnat RNAi (v32255), 5xUAS-dicer2; dWnk$^{3L.282}$ FRT2A 5xUAS-dWnk$^{S632A}$ /tub-gal80 FRT2A |
| C,D 4 | w; OK371-gal4, 10xUAS-ivs-mCD8-GFP, aseFLP$^{2e}$/5xUAS-Nmnat RNAi (v32255), 5xUAS-dicer2; dWnk$^{3L.282}$ FRT2A 5xUAS-dWnk$^{D618A}$ /tub-gal80 FRT2A |
| C,D 5 | w; OK371-gal4, 10xUAS-ivs-mCD8-GFP, aseFLP$^{2e}$/5xUAS-Nmnat RNAi (v32255), 5xUAS-dicer2; dWnk$^{3L.282}$ FRT2A 5xUAS-dWnk$^{L619F}$ /tub-gal80 FRT2A |

Figure 3:

| | |
|---|---|
| A,D,E 1 | w; OK371-gal4, 10xUAS-ivs-mCD8-GFP, aseFLP$^{2c}$/5xUAS-Nmnat RNAi (v32255), 5xUAS-dicer2; FRT2A FRT82B/FRT82B tub-gal80 |
| A,D,E 2 | w; OK371-gal4, 10xUAS-ivs-mCD8-GFP, aseFLP$^{2c}$/5xUAS-Nmnat RNAi (v32255), 5xUAS-dicer2; FRT82B fray$^{07551}$/FRT82B tub-gal80 |
| A,D,E 3 | w 5xUAS-Fray$^{T206E}$/w or y; OK371-gal4, 10xUAS-ivs-mCD8-GFP, aseFLP$^{2c}$/5xUAS-Nmnat RNAi (v32255), 5xUAS-dicer2; FRT82B fray$^{07551}$/FRT82B tub-gal80 |
| A,D,E 4 | w 5xUAS-Fray$^{S347D}$/w or y; OK371-gal4, 10xUAS-ivs-mCD8-GFP, aseFLP$^{2c}$/5xUAS-Nmnat RNAi (v32255), 5xUAS-dicer2; FRT82B fray$^{07551}$/FRT82B tub-gal80 |
| A,D,E 5 | w 5xUAS-Fray$^{D185A}$/w or y; OK371-gal4, 10xUAS-ivs-mCD8-GFP, aseFLP$^{2c}$/5xUAS-Nmnat RNAi (v32255), 5xUAS-dicer2; FRT82B fray$^{07551}$/FRT82B tub-gal80 |
| A,D,E 6 | w 5xUAS-dWnk$^{L619F}$/w or y; OK371-gal4, 10xUAS-ivs-mCD8-GFP, aseFLP$^{2c}$/5xUAS-Nmnat RNAi (v32255), 5xUAS-dicer2; FRT82B fray$^{07551}$/FRT82B tub-gal80 |
| C1 | w; OK371-gal4, 10xUAS-ivs-mCD8-GFP, aseFLP$^{2c}$/5xUAS-Nmnat RNAi (v32255), 5xUAS-dicer2; FRT2A FRT82B/FRT82B tub-gal80 |
| C2 | w; OK371-gal4, 10xUAS-ivs-mCD8-GFP, aseFLP$^{2c}$/5xUAS-Nmnat RNAi (v32255), 5xUAS-dicer2; FRT82B fray$^{07551}$/FRT82B tub-gal80 |
| C3 | w; OK371-gal4, 10xUAS-ivs-mCD8-GFP, aseFLP$^{2c}$/5xUAS-Nmnat RNAi (v32255), 5xUAS-dicer2; FRT82B fray$^{\Delta}$/FRT82B tub-gal80 |
| C4 | w; OK371-gal4, 10xUAS-ivs-mCD8-GFP, aseFLP$^{2c}$/5xUAS-Nmnat RNAi (v32255), 5xUAS-dicer2; FRT82B fray$^{K67M}$/FRT82B tub-gal80 |
| F1 | w; OK371-gal4, 10xUAS-ivs-mCD8-GFP, aseFLP$^{2e}$/5xUAS-Nmnat RNAi (v32255), 5xUAS-dicer2; FRT2A FRT82B/tub-gal80 FRT2A |
| F2 | w; OK371-gal4, 10xUAS-ivs-mCD8-GFP, aseFLP$^{2e}$/5xUAS-Nmnat RNAi (v32255), 5xUAS-dicer2; dWnk$^{3L.282}$ FRT2A FRT82B/tub-gal80 FRT2A |
| F3 | w; OK371-gal4, 10xUAS-ivs-mCD8-GFP, aseFLP$^{2e}$/5xUAS-Nmnat RNAi (v32255), 5xUAS-dicer2; dWnk$^{3L.282}$ FRT2A 5xUAS-Fray$^{S347D}$/tub-gal80 FRT2A |

| | |
|---|---|
| F4 | w; OK371-gal4, 10xUAS-ivs-mCD8-GFP, aseFLP$^{2e}$/5xUAS-Nmnat RNAi (v32255), 5xUAS-dicer2; dWnk$^{3L.282}$ FRT2A 5xUAS-Fray$^{D185A}$/tub-gal80 FRT2A |

Figure 4:

| | |
|---|---|
| A-C 1 | w; OK371-gal4, 10xUAS-ivs-mCD8-GFP, aseFLP$^{2e}$/5xUAS-Nmnat RNAi (v32255), 5xUAS-dicer2; FRT2A FRT82B/tub-gal80 FRT2A |
| A-C 2 | w; OK371-gal4, 10xUAS-ivs-mCD8-GFP, aseFLP$^{2e}$/5xUAS-Nmnat RNAi (v32255), 5xUAS-dicer2; dWnk$^{3L.282}$ FRT2A FRT82B/tub-gal80 FRT2A |
| A-C 3 | w; OK371-gal4, 10xUAS-ivs-mCD8-GFP, aseFLP$^{2e}$/5xUAS-Nmnat RNAi (v32255), 5xUAS-dicer2; dSarm$^{896}$ FRT2A FRT82B/tub-gal80 FRT2A |
| A-C 4 | w; OK371-gal4, 10xUAS-ivs-mCD8-GFP, aseFLP$^{2e}$/5xUAS-Nmnat RNAi (v32255), 5xUAS-dicer2; dSarm$^{896}$ dWnk$^{3L.282}$ FRT2A/tub-gal80 FRT2A |
| A-C 5 | w; OK371-gal4, 10xUAS-ivs-mCD8-GFP, aseFLP$^{2e}$/5xUAS-Nmnat RNAi (v32255), 5xUAS-dicer2; axed$^{3L.11}$ FRT2A FRT82B/tub-gal80 FRT2A |
| D1 | w; OK371-gal4, 10xUAS-ivs-mCD8-GFP, aseFLP$^{2e}$/5xUAS-dSarm$^{\Delta ARM}$-myc, tub-gal80$^{ts}$; FRT2A FRT82B/tub-gal80 FRT2A |
| D2 | w; OK371-gal4, 10xUAS-ivs-mCD8-GFP, aseFLP$^{2e}$/5xUAS-dSarm$^{\Delta ARM}$-myc, tub-gal80$^{ts}$; dWnk$^{3L.282}$ FRT2A FRT82B/tub-gal80 FRT2A |
| D3 | w; OK371-gal4, 10xUAS-ivs-mCD8-GFP, aseFLP$^{2c}$/5xUAS-dSarm$^{\Delta ARM}$-myc, tub-gal80$^{ts}$; FRT2A FRT82B/FRT82B tub-gal80 |
| D4 | w; OK371-gal4, 10xUAS-ivs-mCD8-GFP, aseFLP$^{2c}$/5xUAS-dSarm$^{\Delta ARM}$-myc, tub-gal80$^{ts}$; FRT82B fray$^{07551}$/FRT82B tub-gal80 |
| E,F 1 | w; OK371-gal4, 10xUAS-ivs-mCD8-GFP, aseFLP$^{2e}$/5xUAS-Nmnat RNAi (v32255), 5xUAS-dicer2; FRT2A FRT82B/tub-gal80 FRT2A |
| E,F 2 | w; OK371-gal4, 10xUAS-ivs-mCD8-GFP, aseFLP$^{2e}$/5xUAS-Nmnat RNAi (v32255), 5xUAS-dicer2; dSarm$^{896}$ FRT2A FRT82B/tub-gal80 FRT2A |
| E,F 3 | w; OK371-gal4, 10xUAS-ivs-mCD8-GFP, aseFLP$^{2e}$/5xUAS-Nmnat RNAi (v32255), 5xUAS-dicer2; dSarm$^{896}$ FRT2A 5xUAS-dWnk$^{L619F}$/tub-gal80 FRT2A |
| E,F 4 | w; OK371-gal4, 10xUAS-ivs-mCD8-GFP, aseFLP$^{2e}$/5xUAS-Nmnat RNAi (v32255), 5xUAS-dicer2; dSarm$^{896}$ FRT2A 5xUAS-dWnk$^{S632A}$/tub-gal80 FRT2A |
| G,H 1 | w; OK371-gal4, 10xUAS-ivs-mCD8-GFP, aseFLP$^{2e}$/5xUAS-Nmnat RNAi (v32255), 5xUAS-dicer2; FRT2A FRT82B/tub-gal80 FRT2A |
| G,H 2 | w; OK371-gal4, 10xUAS-ivs-mCD8-GFP, aseFLP$^{2e}$/5xUAS-Nmnat RNAi (v32255), 5xUAS-dicer2; dSarm$^{896}$ FRT2A FRT82B/tub-gal80 FRT2A |
| G,H 3 | w; OK371-gal4, 10xUAS-ivs-mCD8-GFP, aseFLP$^{2e}$/5xUAS-Nmnat RNAi (v32255), 5xUAS-dicer2; dSarm$^{896}$ FRT2A 5xUAS-Fray$^{S347D}$/tub-gal80 FRT2A |
| G,H 4 | w; OK371-gal4, 10xUAS-ivs-mCD8-GFP, aseFLP$^{2e}$/5xUAS-Nmnat RNAi (v32255), 5xUAS-dicer2; dSarm$^{896}$ FRT2A 5xUAS-Fray$^{D185A}$/tub-gal80 FRT2A |
| I1 | w; OK371-gal4, 10xUAS-ivs-mCD8-GFP, aseFLP$^{2e}$/sp; FRT2A, 5xUAS-LacZ/tub-gal80 FRT2A |
| I2 | w; OK371-gal4, 10xUAS-ivs-mCD8-GFP, aseFLP$^{2e}$/Sp; FRT2A 5xUAS-dWnk$^{L619F}$/tub-gal80 FRT2A |
| I3 | w; OK371-gal4, 10xUAS-ivs-mCD8-GFP, aseFLP$^{2e}$/Sp; FRT2A 5xUAS-dWnk$^{S632A}$/tub-gal80 FRT2A |
| I4 | w; OK371-gal4, 10xUAS-ivs-mCD8-GFP, aseFLP$^{2e}$/Sp; FRT2A 5xUAS-Fray$^{S347D}$/tub-gal80 FRT2A |
| I5 | w; OK371-gal4, 10xUAS-ivs-mCD8-GFP, aseFLP$^{2e}$/Sp; FRT2A 5xUAS-Fray$^{D185A}$/tub-gal80 FRT2A |

Figure 5:

| | |
|---|---|
| A1 | w; OK371-gal4, 10xUAS-ivs-mCD8-GFP, aseFLP$^{2e}$/5xUAS-Nmnat RNAi (v32255), 5xUAS-dicer2; FRT2A FRT82B/tub-gal80 FRT2A |
| A2 | w; OK371-gal4, 10xUAS-ivs-mCD8-GFP, aseFLP$^{2e}$/5xUAS-Nmnat RNAi (v32255), 5xUAS-dicer2; axed$^{3L.11}$ FRT2A FRT82B/tub-gal80 FRT2A |
| A3 | w; OK371-gal4, 10xUAS-ivs-mCD8-GFP, aseFLP$^{2e}$/5xUAS-Nmnat RNAi (v32255), 5xUAS-dicer2; axed$^{3L.11}$ FRT2A 5xUAS-Fray$^{S347D}$ /tub-gal80 FRT2A |
| C1 | w; OK371-gal4, 10xUAS-ivs-mCD8-GFP, aseFLP$^{2e}$/Sp; FRT2A FRT82B/tub-gal80 FRT2A |
| C2 | w; OK371-gal4, 10xUAS-ivs-mCD8-GFP, aseFLP$^{2e}$/Sp; dWnk$^{3L.282}$ FRT2A FRT82B/tub-gal80 FRT2A |
| C3 | w; OK371-gal4, 10xUAS-ivs-mCD8-GFP, aseFLP$^{2e}$/Sp; dSarm$^{E1170A}$ FRT2A/tub-gal80 FRT2A |
| C4 | w; OK371-gal4, 10xUAS-ivs-mCD8-GFP, aseFLP$^{2e}$/Sp; dSarm$^{E1170A}$ dWnk$^{3L.282}$ FRT2A/tub-gal80 FRT2A |
| C5 | w; OK371-gal4, 10xUAS-ivs-mCD8-GFP, aseFLP$^{2e}$/Sp; dSarm$^{896}$ FRT2A FRT82B/tub-gal80 FRT2A |
| C6 | w; OK371-gal4, 10xUAS-ivs-mCD8-GFP, aseFLP$^{2e}$/Sp; dSarm$^{896}$ FRT2A 5xUAS-Fray$^{S347D}$/tub-gal80 FRT2A |

Figure S1:

| | |
|---|---|
| D,E | *w; OK371-gal4, 10xUAS-ivs-mCD8-GFP, aseFLP*$^{2e}$*/5xUAS-Nmnat RNAi (v32255), 5xUAS-dicer2; FRT2A FRT82B/tub-gal80 FRT2A* |
| F1 | *w; OK371-gal4, 10xUAS-ivs-mCD8-GFP, aseFLP*$^{2e}$*/5xUAS-Nmnat RNAi (v32255), 5xUAS-dicer2; FRT2A FRT82B/tub-gal80 FRT2A* |
| F2 | *w, 5xUAS-dWnk*$^{WT}$*/w or y; OK371-gal4, 10xUAS-ivs-mCD8-GFP, aseFLP*$^{2e}$*/5xUAS-Nmnat RNAi (v32255), 5xUAS-dicer2; FRT2A FRT82B/tub-gal80 FRT2A* |
| F3 | *w; OK371-gal4, 10xUAS-ivs-mCD8-GFP, aseFLP*$^{2e}$*/5xUAS-Nmnat RNAi (v32255), 5xUAS-dicer2; dWnk*$^{3L.1541}$ *FRT2A FRT82B/tub-gal80 FRT2A* |
| F4 | *w, 5xUAS-dWnk*$^{WT}$*/w or y; OK371-gal4, 10xUAS-ivs-mCD8-GFP, aseFLP*$^{2e}$*/5xUAS-Nmnat RNAi (v32255), 5xUAS-dicer2; dWnk*$^{3L.1541}$ *FRT2A FRT82B/tub-gal80 FRT2A* |
| F5 | *w; OK371-gal4, 10xUAS-ivs-myrTdTom, aseFLP*$^{2c}$*/5xUAS-Nmnat RNAi (v32255), 5xUAS-dicer2; dWnk*$^{MB06499}$ *FRT2A/tub-gal80 FRT2A* |
| F6 | *w, 5xUAS-dWnk*$^{WT}$*/w or y; OK371-gal4, 10xUAS-ivs-myrTdTom, aseFLP*$^{2c}$*/5xUAS-Nmnat RNAi (v32255), 5xUAS-dicer2; dWnk*$^{MB0649}$*FRT2A/tub-gal80 FRT2A* |
| F7 | *w; OK371-gal4, 10xUAS-ivs-mCD8-GFP, aseFLP*$^{2e}$*/5xUAS-Nmnat RNAi (v32255), 5xUAS-dicer2; Df(3L)Exel9066 FRT2A/tub-gal80 FRT2A* |
| F8 | *w, 5xUAS-dWnk*$^{WT}$*/w or y; OK371-gal4, 10xUAS-ivs-mCD8-GFP, aseFLP*$^{2e}$*/5xUAS-Nmnat RNAi (v32255), 5xUAS-dicer2; Df(3L)Exel9066 FRT2A/tub-gal80 FRT2A* |
| G1 | *w; OK371-gal4, 10xUAS-ivs-mCD8-GFP, aseFLP*$^{2e}$*/5xUAS-Nmnat RNAi (v32255), 5xUAS-dicer2; FRT80B/tub-gal80 FRT80B* |
| G2 | *w; OK371-gal4, 10xUAS-ivs-mCD8-GFP, aseFLP*$^{2e}$*/5xUAS-Nmnat RNAi (v32255), 5xUAS-dicer2; dWnk*$^{F1183}$ *FRT80B/tub-gal80 FRT80B* |
| G3 | *w; OK371-gal4, 10xUAS-ivs-mCD8-GFP, aseFLP*$^{2e}$*/5xUAS-Nmnat RNAi (v32255), 5xUAS-dicer2; dWnk*$^{G1286}$ *FRT80B/tub-gal80 FRT80B* |
| H1 | *w; OK371-gal4, 10xUAS-ivs-mCD8-GFP, aseFLP*$^{2e}$*/Sp; FRT2A FRT82B/tub-gal80 FRT2A* |
| H2 | *w; OK371-gal4, 10xUAS-ivs-mCD8-GFP, aseFLP*$^{2e}$*/Sp; dWnk*$^{F1183}$ *FRT80B/tub-gal80 FRT80B* |
| H3 | *w; OK371-gal4, 10xUAS-ivs-mCD8-GFP, aseFLP*$^{2e}$*/Sp; dWnk*$^{G1286}$ *FRT80B/tub-gal80 FRT80B* |
| H4 | *w; OK371-gal4, 10xUAS-ivs-mCD8-GFP, aseFLP*$^{2e}$*/Sp; dWnk*$^{3L.282}$ *FRT2A FRT82B/tub-gal80 FRT2A* |
| H5 | *w; OK371-gal4, 10xUAS-ivs-myrTdTom, aseFLP*$^{2c}$*/Sp; dWnk*$^{MB06499}$ *FRT2A/tub-gal80 FRT2A* |
| H6 | *w; OK371-gal4, 10xUAS-ivs-mCD8-GFP, aseFLP*$^{2e}$*/Sp; Df(3L)Exel9066 FRT2A/tub-gal80 FRT2A* |
| I1 | *w; OK371-gal4, 10xUAS-ivs-mCD8-GFP, aseFLP*$^{2e}$*/Sp; FRT2A FRT82B/tub-gal80 FRT2A* |
| I2 | *w, 5xUAS-dWnk*$^{WT}$*/w or y; OK371-gal4, 10xUAS-ivs-mCD8-GFP, aseFLP*$^{2e}$*/Sp; FRT2A FRT82B/tub-gal80 FRT2A* |

Figure S2:

| | |
|---|---|
| A1 | *w; OK371-gal4, 10xUAS-ivs-mCD8-GFP, aseFLP*$^{2e}$*/Sp; FRT2A, 5xUAS-LacZ/tub-gal80 FRT2A* |
| A2 | *w; OK371-gal4, 10xUAS-ivs-mCD8-GFP, aseFLP*$^{2e}$*/Sp; FRT2A 5xUAS-dWnk*$^{S632A}$*/tub-gal80 FRT2A* |
| A3 | *w; OK371-gal4, 10xUAS-ivs-mCD8-GFP, aseFLP*$^{2e}$*/Sp; FRT2A 5xUAS-dWnk*$^{D618A}$*/tub-gal80 FRT2A* |
| A4 | *w; OK371-gal4, 10xUAS-ivs-mCD8-GFP, aseFLP*$^{2e}$*/Sp; FRT2A 5xUAS-dWnk*$^{L619F}$*/tub-gal80 FRT2A* |
| B1-4 | Like Fig. 2, B1-4 |
| C,E 1-5 | Like Fig. 2, C,D 1-5 |
| F,G 1 | *w; OK371-gal4, 10xUAS-ivs-mCD8-GFP, aseFLP*$^{2e}$*/5xUAS-Nmnat RNAi (v32255), 5xUAS-dicer2; FRT2A FRT82B/tub-gal80 FRT2A* |
| F,G 2 | *w; OK371-gal4, 10xUAS-ivs-mCD8-GFP, aseFLP*$^{2e}$*/5xUAS-Nmnat RNAi (v32255), 5xUAS-dicer2; Df(3L)Exel9066 FRT2A/tub-gal80 FRT2A* |
| F,G 3 | *w; OK371-gal4, 10xUAS-ivs-mCD8-GFP, aseFLP*$^{2e}$*/5xUAS-Nmnat RNAi (v32255), 5xUAS-dicer2; Df(3L)Exel9066 FRT2A 5xUAS-dWnk*$^{S632A}$*/tub-gal80 FRT2A* |
| F,G 4 | *w; OK371-gal4, 10xUAS-ivs-myrTdTom, aseFLP*$^{2c}$*/5xUAS-Nmnat RNAi (v32255), 5xUAS-dicer2; dWnk*$^{MB06499}$ *FRT2A/tub-gal80 FRT2A* |
| F,G 5 | *w; OK371-gal4, 10xUAS-ivs-myrTdTom, aseFLP*$^{2c}$*/5xUAS-Nmnat RNAi (v32255), 5xUAS-dicer2; dWnk*$^{MB06499}$ *FRT2A 5xUAS-dWnk*$^{S632A}$ */tub-gal80 FRT2A* |

Figure S3:

| | |
|---|---|
| C-E 1-4 | Like Fig. 3F 1-4 |
| F,G 1 | *w; OK371-gal4, 10xUAS-ivs-myrTdTom, aseFLP*$^{2c}$*/5xUAS-Nmnat RNAi (v32255), 5xUAS-dicer2; FRT2A FRT82B/tub-gal80 FRT2A* |

| | |
|---|---|
| F,G 2 | *w; OK371-gal4, 10xUAS-ivs-myrTdTom, aseFLP$^{2c}$/5xUAS-Nmnat RNAi (v32255), 5xUAS-dicer2; dWnk$^{MB06499}$ FRT2A/tub-gal80 FRT2A* |
| F,G 3 | *w; OK371-gal4, 10xUAS-ivs-myrTdTom, aseFLP$^{2c}$/5xUAS-Nmnat RNAi (v32255), 5xUAS-dicer2; dWnk$^{MB06499}$ FRT2A 5xUAS-Fray$^{S347D}$/tub-gal80 FRT2A* |
| F,G 4 | *w; OK371-gal4, 10xUAS-ivs-myrTdTom, aseFLP$^{2c}$/5xUAS-Nmnat RNAi (v32255), 5xUAS-dicer2; dWnk$^{MB06499}$ FRT2A 5xUAS-Fray$^{D185A}$/tub-gal80 FRT2A* |
| H1 | *w; OK371-gal4, 10xUAS-ivs-mCD8-GFP, aseFLP$^{2e}$/Sp; FRT2A FRT82B/tub-gal80 FRT2A* |
| H2 | *w; OK371-gal4, 10xUAS-ivs-mCD8-GFP, aseFLP$^{2e}$/Sp; FRT2A 5xUAS-Fray$^{T206E}$/tub-gal80 FRT2A* |
| H3 | *w; OK371-gal4, 10xUAS-ivs-mCD8-GFP, aseFLP$^{2e}$/Sp; FRT2A 5xUAS-Fray$^{S347D}$/tub-gal80 FRT2A* |
| H4 | *w; OK371-gal4, 10xUAS-ivs-mCD8-GFP, aseFLP$^{2e}$/Sp; FRT2A 5xUAS-Fray$^{D185A}$/tub-gal80 FRT2A* |
| H5 | *w; OK371-gal4, 10xUAS-ivs-mCD8-GFP, aseFLP$^{2e}$/Sp; FRT2A 5xUAS-dWnk$^{L619F}$/tub-gal80 FRT2A* |

Figure S4:

| | |
|---|---|
| A-B 1-5 | Like Fig. 4A-C 1-5 |
| C,D 1 | *w; OK371-gal4, 10xUAS-ivs-myrTdTom, aseFLP$^{2c}$/5xUAS-Nmnat RNAi (v32255), 5xUAS-dicer2; FRT2A FRT82B/tub-gal80 FRT2A* |
| C,D 2 | *w; OK371-gal4, 10xUAS-ivs-myrTdTom, aseFLP$^{2c}$/5xUAS-Nmnat RNAi (v32255), 5xUAS-dicer2; dWnk$^{MB06499}$ FRT2A/tub-gal80 FRT2A* |
| C,D 3 | *w; OK371-gal4, 10xUAS-ivs-myrTdTom, aseFLP$^{2c}$/5xUAS-Nmnat RNAi (v32255), 5xUAS-dicer2; dSarm$^{896}$ FRT2A FRT82B/tub-gal80 FRT2A* |
| C,D 4 | *w; OK371-gal4, 10xUAS-ivs-myrTdTom, aseFLP$^{2c}$/5xUAS-Nmnat RNAi (v32255), 5xUAS-dicer2; dSarm$^{896}$ dWnk$^{MB06499}$ FRT2A/tub-gal80 FRT2A* |
| C,D 5 | *w; OK371-gal4, 10xUAS-ivs-myrTdTom, aseFLP$^{2c}$/5xUAS-Nmnat RNAi (v32255), 5xUAS-dicer2; axed$^{3L.11}$ FRT2A FRT82B/tub-gal80 FRT2A* |

Figure S5:

| | |
|---|---|
| A,B 1 | *w; OK371-gal4, 10xUAS-ivs-mCD8-GFP, aseFLP$^{2e}$/5xUAS-Nmnat RNAi (v32255), 5xUAS-dicer2; FRT2A FRT82B/tub-gal80 FRT2A* |
| A,B 2 | *w; OK371-gal4, 10xUAS-ivs-mCD8-GFP, aseFLP$^{2e}$/5xUAS-Nmnat RNAi (v32255), 5xUAS-dicer2; Ncc69$^{r2}$ FRT2A/tub-gal80 FRT2A* |
| A,B 3 | *w; OK371-gal4, 10xUAS-ivs-myrTdTom, aseFLP$^{2c}$/5xUAS-Nmnat RNAi (v32255), 5xUAS-dicer2; FRT2A FRT82B/FRT82B tub-gal80* |
| A,B 4 | *w; OK371-gal4, 10xUAS-ivs-myrTdTom, aseFLP$^{2c}$/5xUAS-Nmnat RNAi (v32255), 5xUAS-dicer2; FRT2A FRT82B Irk2$^{CR70959-TG4.2}$/FRT82B tub-gal80* |
| A,B 5 | *w; OK371-gal4, 10xUAS-ivs-myrTdTom, aseFLP$^{2c}$/5xUAS-Nmnat RNAi (v32255), 5xUAS-dicer2; FRT2A FRT82B Irk1$^{MI08404}$/FRT82B tub-gal80* |
| A,B 6 | *w; FRT40A FRTG13/tub-gal80 FRT40A; elaV-gal4, 10xUAS-ivs-mCD8-GFP, aseFLP$^{3b}$/20xUAS-Nmnat RNAi, 5xUAS-dicer2* |
| A,B 7 | *w; Irk3$^{SK1}$ FRT40A/tub-gal80 FRT40A; elaV-gal4, 10xUAS-ivs-mCD8-GFP, aseFLP$^{3b}$/20xUAS-Nmnat RNAi, 5xUAS-dicer2* |
